# A unified framework for measuring selection on cellular lineages and traits

**DOI:** 10.1101/2021.07.18.452747

**Authors:** Shunpei Yamauchi, Takashi Nozoe, Reiko Okura, Edo Kussell, Yuichi Wakamoto

## Abstract

Intracellular states probed by gene expression profiles and metabolic activities are intrinsically noisy, causing phenotypic variations among cellular lineages. Understanding the adaptive and evolutionary roles of such variations requires clarifying their linkage to population growth rates. Extending a cell lineage statistics framework, here we show that a population’s growth rate can be expanded by fitness cumulants of any cell lineage trait. The expansion enables quantifying the contribution of each fitness cumulant, such as variance and skewness, to population growth. We introduce a function that contains all the essential information of cell lineage statistics, including mean lineage fitness and selection strength. We reveal a relation between fitness heterogeneity and population growth rate response to perturbation. We apply the framework to experimental cell lineage data from bacteria to mammalian cells, revealing that third or higher-order cumulants’ contributions are negligible under constant growth conditions but could be significant in regrowing processes from growth-arrested conditions. Furthermore, we identify cellular populations in which selection leads to an increase of fitness variance among lineages. The framework assumes no particular growth models or environmental conditions, and is thus applicable to various biological phenomena for which phenotypic heterogeneity and cellular proliferation are important.

## Introduction

Growth rates of cellular populations are physiological quantities directly linked to the fitness of cellular organisms. To understand the roles of biological processes and reactions within cells, including modulation of gene expression and metabolic states, one must characterize how they are eventually channeled into an increase or maintenance of population growth rates.

As documented by many single-cell studies, phenotypic states of individual cells in cellular populations are heterogeneous and often correlate with fitness variations among cellular lineages [1–8]. Fitness heterogeneity within a population causes a statistical bias on ancestral cells’ contributions to the number of descendants, which is broadly referred to as “selection” [9]. Such bias from growth heterogeneity makes the relations between cellular lineages and populations nontrivial. For example, an intriguing consequence of intra-population selection is a growth rate gain, a phenomenon that cell population’s growth rate becomes greater than the mean division rate of isolated single-cell lineages [4, 10, 11]. Recent progress of single-cell measurements has enabled high-throughput acquisitions of cellular lineage trees and historical dynamics in each lineage [6, 10, 12]. However, establishing the theory and method of cellular lineage statistics to quantify fitness differences among different phenotypic states and intrapopulation selection is still in progress [13–16].

Growth of cellular populations can be described using the ensemble of individual cells’ growth histories [9]. A theoretical approach that regards a cell lineage (history) as a basic unit of analysis has offered illuminating insights into population dynamics. For example, it has provided the formula for untangling selection from responses [9], population response to age-specific changes in mortality and fecundity [17], fluctuation relations of fitness [16, 18], and relations between cell size growth rate and population growth rate [19, 20].

Employing this cell history-based formulation of population dynamics, we have previously proposed a method of cellular lineage statistics that allows quantification of fitness landscapes and selection strength for any traits of cellular lineages [13]. Here, we extend this statistical framework and show that population growth rates can be expanded by fitness cumulants of cellular lineage traits. The expansion allows us to quantify the contributions of fitness cumulants, such as variance and skewness, to overall population growth rates. We derive a relation for the population growth rate’s response to perturbation, and define a function that provides all the essential information of lineage statistics. We apply the framework to experimental single-cell lineage data of bacteria, yeast, and mammalian cells. We show that contributions of third or higher-order fitness cumulants to population growth rates are negligible in most constantly growing populations but can be significant under environmental shift conditions. We measure the fitness landscapes for a growth-regulating sigma factor in *E. coli* and identify the conditions where its continuum expression heterogeneity correlates with lineage fitness in cellular populations.

### Theoretical background

First, we briefly review the analytical framework of cell lineage statistics introduced in [13]. This framework allows us to quantitatively infer fitness differences associated with distinct states of cellular lineage traits and selection within a growing cell population from empirical single-cell lineage tree data. Time-lapse single-cell measurements provide cellular growth and division information in the form of lineage trees (Fig. 1) [12]. We regard a lineage *σ* as a cell history traceable back from a descendant cell at the final time point *t* = *τ* (Fig. 1B). For the case of cellular growth shown in Fig. 1A, 22 cell lineages exist in the trees.

**Figure 1.**
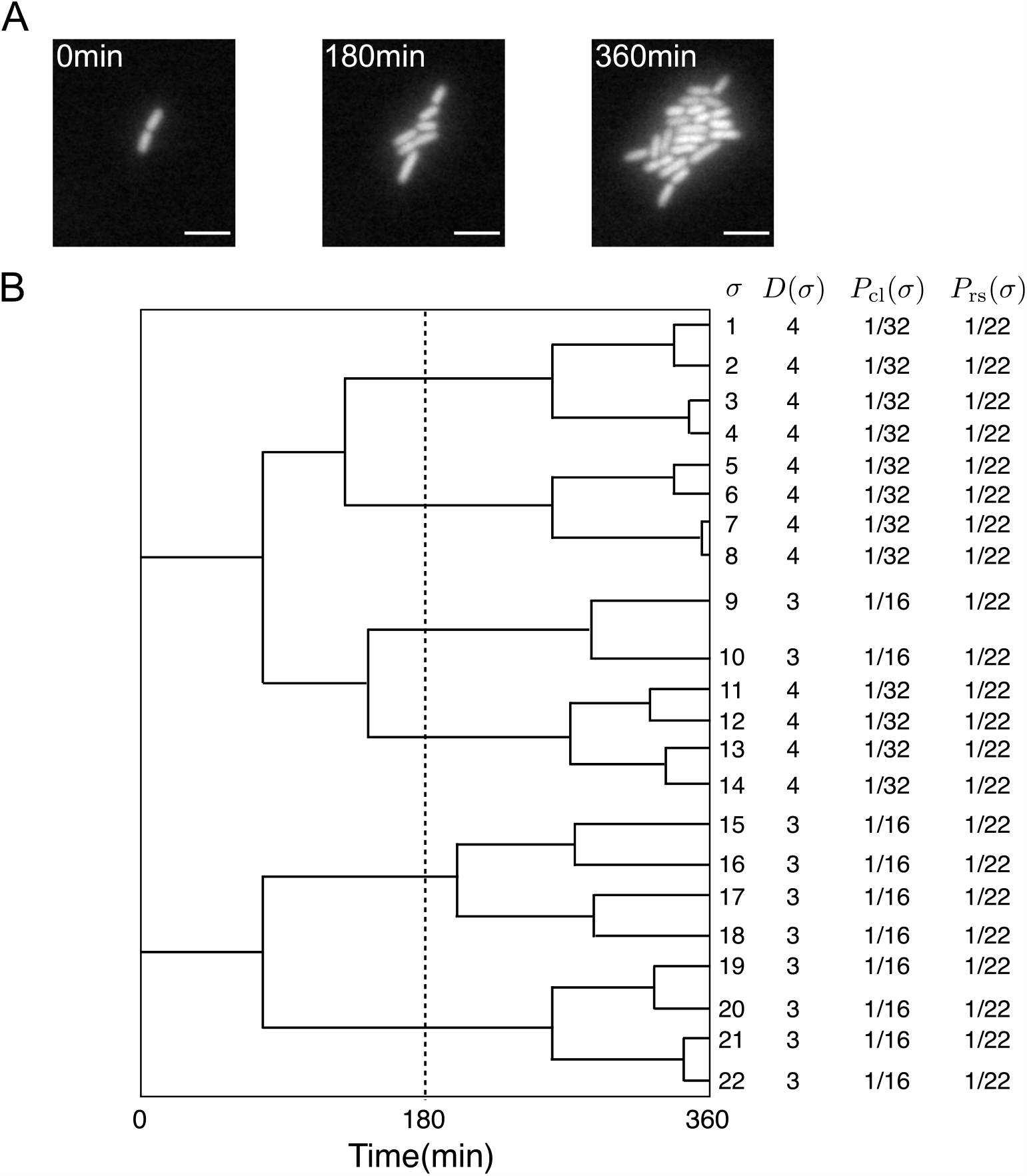
Representative single-cell lineage trees. **A**. Time-lapse images of a growing microcolony of *Escherichia coli* expressing green fluorescent protein (GFP) from plasmids. Scale bars, 5 *µ*m. **B**. Cellular lineage trees for the microcolony in A. Bifurcations in the trees represent cell divisions. *σ* denotes cell lineage labels. *D*(*σ*) shows the number of cell divisions in each lineage. *P*_cl_(*σ*) and *P*_rs_(*σ*) are chronological and retrospective probabilities defined in the main text.

We assign two types of probability weight to cellular lineages. One is retrospective probability, in which we assign equal weight *P*_rs_(*σ*) := 1*/N*_*τ*_ to all lineages, where *N*_*τ*_ is the number of cells at the final time point *t* = *τ*. *P*_rs_(*σ*) represents the probability of selecting the history of a cell present at the endpoints of lineage trees. Another is chronological probability, in which we assign the weight *P*_cl_(*σ*) := 2^−*D*(*σ*)^*/N*_0_ to the lineages, where *D*(*σ*) is the number of cell divisions on lineage *σ* and *N*_0_ is the initial number of cells at *t* = 0. *P*_cl_(*σ*) represents the probability of choosing lineage *σ* descending the tree from one of the ancestor cells at *t* = 0 and selecting one branch with the probability 1*/*2 at every cell division. *P*_rs_(*σ*) and *P*_cl_(*σ*) can be different in general when the number of cell divisions are variable among the cell lineages, as shown in Fig. 1B.

These two types of lineage probability satisfy

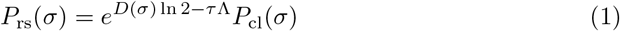

where 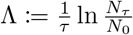 is the population growth rate, and *D*(*σ*) ln 2 is the lineage fitness. This relation shows that cellular lineages with division rates (*D*(*σ*) ln 2*/τ*) higher than population growth rate are overrepresented in the retrospective probability compared to the chronological probability. Summing both sides of (Eq. 1) yields

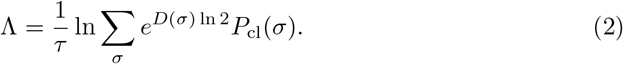

We also define retrospective and chronological probabilities for a *lineage trait X* as 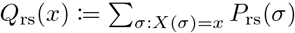 and 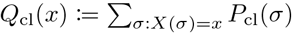, where *X*(*σ*) is the value of trait *X* for lineage *σ*. Here, we regard any measurable quantity associated with cellular lineages as a lineage trait *X*. For example, time-averaged expression levels and production rates of a drug-resistance protein were analyzed as lineage traits in the experiments of [13]. *D* is also a lineage trait and has exceptional significance as its selection strength defined below sets the maximum bound for the selection strength of any lineage trait (Supplemental Information).

The *fitness landscape* for lineage trait *X* is defined as

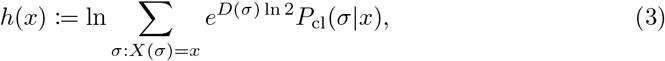

where *P*_cl_(*σ*|*x*) = *P*_cl_(*σ*)*/Q*_cl_(*x*). This definition states that *h*(*x*) represents the net fitness of the subset of lineages with trait value *X* = *x*. From (Eq. 1) and the definitions of *Q*_rs_(*x*) and *h*(*x*), one finds

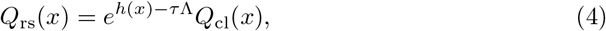

which has the same form as (Eq. 1). The fitness landscape *h*(*x*) is, thus, a natural definition of lineage fitness for a lineage trait *X*. We also remark that by evaluating (Eq. 3) for the trait *X* = *D*, the fitness landscape for trait value *D* = *d* is given by

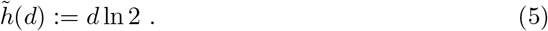

By summing both sides of (Eq. 4) over *x*, we can express the population growth rate in terms of the fitness landscape as

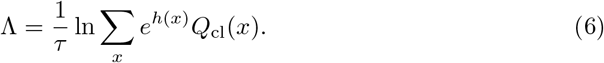

According to (Eq. 4), the trait value *x* is overrepresented in retrospective versus chronological probability when *h*(*x*) *> τ* Λ. Therefore, dissimilarities between *Q*_rs_(*x*) and *Q*_cl_(*x*) can be regarded as measures of selection among different lineage trait values *x*. Following this idea, we define a measure of *selection strength* on trait *X* as

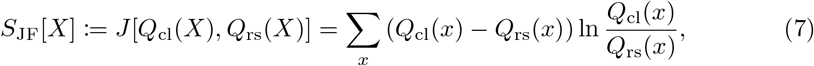

where *J* [*Q*_cl_(*X*), *Q*_rs_(*X*)] is Jefferey’s divergence. From (Eq. 4), we obtain another expression of *S*_JF_[*X*] as

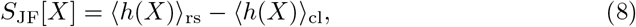

where 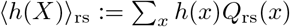 and 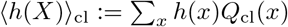 are the retrospective and chronological mean fitness for lineage trait *X*.

Likewise, quantifying the dissimilarities between *Q*_rs_(*x*) and *Q*_cl_(*x*) using the Kullback-Leibler (KL) divergence,

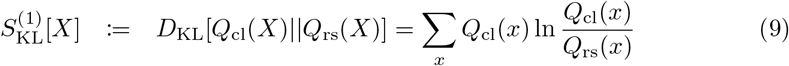

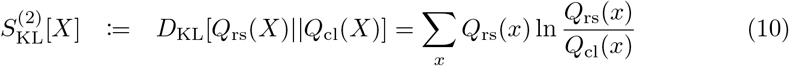

yields two alternative selection strength measures whose sum is *S*_JF_[*X*]. These are also linked to differences in fitness measures:

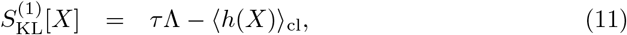

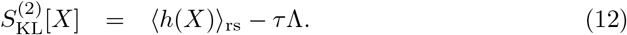

From the non-negativity of the KL-divergence, ⟨*h*(*X*)⟩_rs_ ≥ *τ* Λ ≥ ⟨*h*(*X*) ⟩_cl_. Rewriting (Eq. 11), we obtain

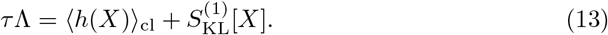

This shows that population growth rates can be decomposed into chronological mean fitness and selection strength. When 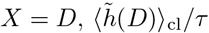 is the mean division rate of cellular lineages without selection.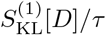, thus, represents growth rate increase by selection compared to the intrinsic division rate of cells. Moreover, 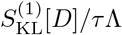 can be regarded as a selection contribution to the population growth rate. Another important property of *D* is that its selection strength sets the maximum bound for the selection strength of any lineage trait, i.e., 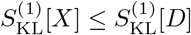 (see Supplemental Information).

Therefore, the quantity

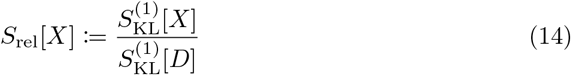

is bounded by 0 ≤*S*_rel_[*X*] ≤1 and evaluates how strongly the heterogeneity of *X* correlates with the division count heterogeneity in a cellular population.

All of the quantities introduced above are measurable without relying on any growth models. Thus, this cell lineage statistics framework is applicable to various single-cell lineage data.

## Results

### Cumulant expansion of population growth rate

To examine the connections between the disparate selection measures and elucidate their meaning, we define a function of a variable *ξ* as

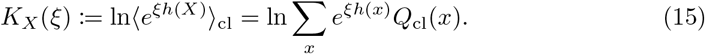

This is the cumulant generating function (cgf) of *h*(*x*) with respect to the chronological distribution *Q*_cl_. We have *K*_*X*_ (0) = 0, and using (Eq. 6) we find

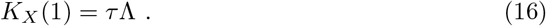

When the radius of covergence of the Taylor expansion of *K*_*X*_ (*ξ*) around *ξ* = 0 is at least 1, *K*_*X*_ (1) can be expressed as the series using the fitness cumulants as

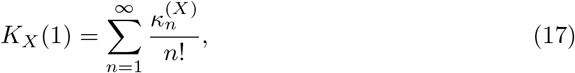

where 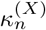 is the *n*-th order fitness cumulant, satisfying 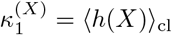, and 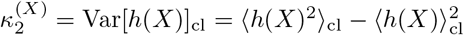. From (Eq. 16) and (Eq. 17),

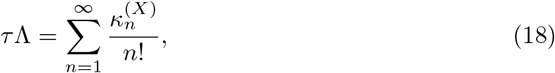

which shows that population growth rates can be expanded by the fitness cumulants of any lineage trait *X*. Additionally, since 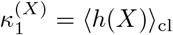, comparing (Eq. 13) and (Eq. 18) yields

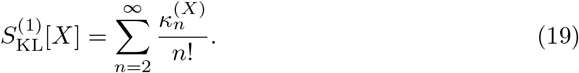

Therefore, 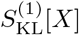 represents the total contribution of second and higher-order fitness cumulants to population growth. When third or higher-order cumulants are negligible, 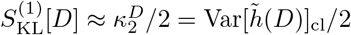 and reflects the contribution of fitness variance to population growth (Supplemental Information).

The cumulant expansion allows us to quantify the relative contributions of various statistical features of fitness distributions to population growth, such as mean, variance, and skewness. We define the cumulative contribution up to the *n*-th order cumulant as

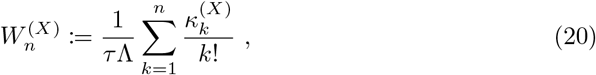

and note that 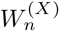 converges to 1 as *n* → ∞.

Since *K*_*X*_ (*ξ*) defined in (Eq. 15) is the cgf of *h*(*x*) with respect to *Q*_cl_(*x*), it provides various forms of fitness and selection measures by simple algebraic calculation, as shown in Table 1. For example, *K*_*X*_(1) = *τ*Λ, 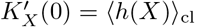, and 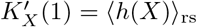.

**Table 1.**
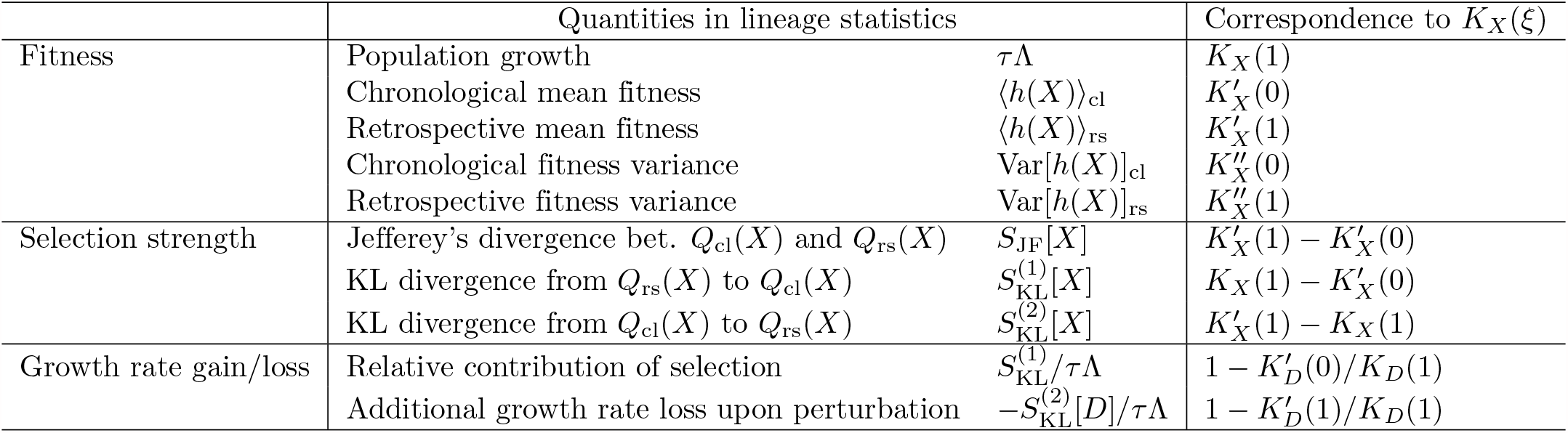
Relationships between *K*_*X*_ (*ξ*) and quantities in cellular lineage statistics.

Therefore, 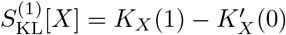.

In general, the *n*-th derivative of *K*_*X*_ (*ξ*) gives the *n*-th order fitness cumulant of the chronological distribution *Q*_cl_(*x*) when evaluated at *ξ* = 0, and of the retrospective distribution *Q*_rs_(*x*) when evaluated at *ξ* = 1. For example, 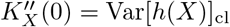 and 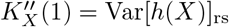 (Supplemental Information). Therefore, *K*_*X*_ (*ξ*) contains complete information on the fitness distributions in both chronological and retrospective statistics.

The relations among the fitness and selection strength measures can be graphically depicted by plotting 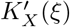 in the interval 0 ≤ *ξ* ≤ 1 (Fig. 2A). For example, the area between the horizontal axis and 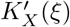 corresponds to population growth rate since 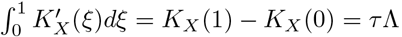. Furthermore, the rectangle between *y* = 0 and 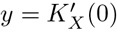 represents chronological mean fitness ⟨*h*(*X*)⟩_cl_, and that between *y* = 0 and 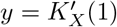 represents retrospective mean fitness ⟨*h*(*X*)⟩_rs_. The selection strength measures, 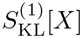 and 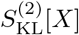 also correspond to the areas shown in Fig. 2A. This graphical representation allows us to capture the contribution of intrapopulation growth heterogeneity to population growth and the difference between 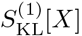 and 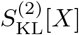 intuitively.

**Figure 2.**
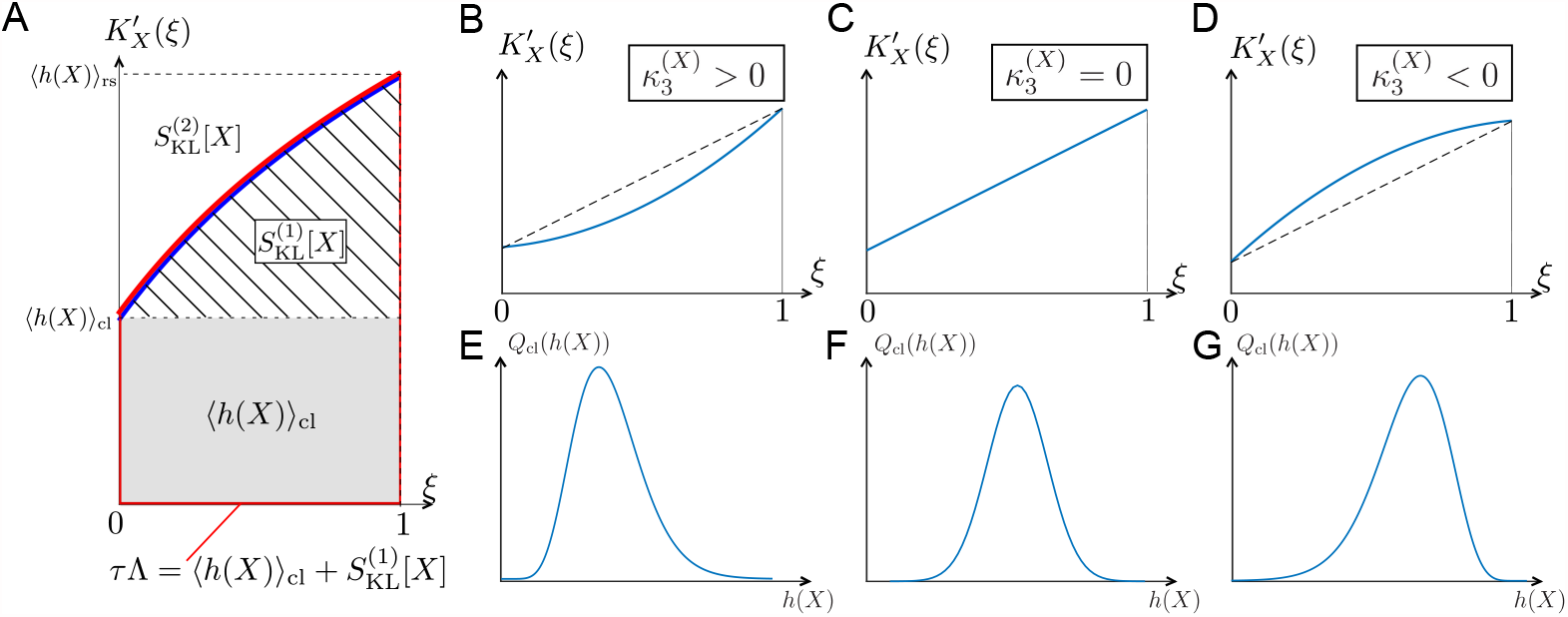
Relationships among chronological distributions’ shape and selection strength measures. **A**. Graphical representation of various fitness and selection strength measures by 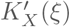-plot. Blue curve represents 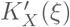. The area between the horizontal axis and 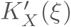 in the interval 0 ≤ *ξ* ≤ 1 outlined in red corresponds to population growth *τ* Λ. The gray and hatched regions correspond to ⟨*h*(*X*)⟩_cl_ and 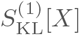, respectively. The area between 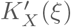 and *y* = ⟨*h*(*X*)⟩_rs_ corresponds to 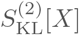. **B-D**. Representative shapes of 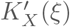 depending on 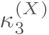. Assuming that the contributions from fourth or higher-order cumulants are negligible, 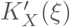 becomes convex downward when 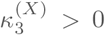 (B); a straight line when 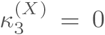 (C); and convex upward when 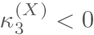 (D). **E-G**. Relationships between third-order fitness cumulant and skewness of chronological distribution *Q*_cl_(*h*).

The difference between 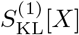 and 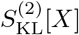 is determined by the higher-order fitness cumulants by the relation

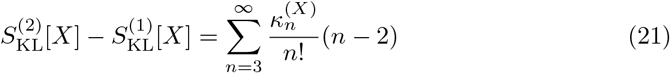

(Supplemental Information). When fourth or higher-order cumulants are negligible, the third-order fitness cumulant 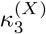, i.e. the skewness of fitness distribution *Q*_cl_(*h*), determines the convexity of 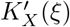 and which selection strength measure is greater, as shown in Fig. 2B-G. Furthermore, when *X* = *D*, fitness landscape is 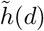 linear with respect to *d* (Eq. 5), and the skewness of division count distribution primarily determines which selection strength is greater.

The slope of the tangent lines to 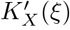 at *ξ* = 0 and 1 corresponds to the chronological and retrospective fitness variances, respectively (Table 1). Therefore, when 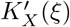 is convex upward in the interval 0 ≤ *ξ* ≤ 1 (as in Fig. 2D), the effect of selection is to decrease the lineage fitness variance, whereas if 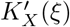 is convex downward (Fig. 2B), selection increases the fitness variance. We indeed find cases of both kinds of behavior in the experimental lineage data, as will be seen below.

### Analytical calculations of fitness measures, selection strength, and fitness cumulants

To observe how the framework works, we show the exact form of *K*_*D*_(*ξ*) for a class of discrete probability distributions containing Poisson, binomial and negative binomial distributions. Let 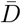 and 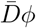 denote the mean and the variance of *Q*_cl_(*D*) respectively (i.e., *ϕ* is the Fano factor of division counts). When *Q*_cl_(*D*) is Poisson, binomial or negative binomial distributions, 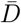 and *ϕ* uniquely determine the form of probability distribution: *ϕ* = 1 for Poisson; *ϕ <* 1 for binomial; and *ϕ >* 1 for negative binomial (Fig. 3A). Then, *K*_*D*_(*ξ*) for these distributions have a closed form

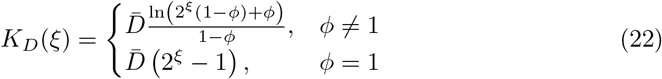

(Supplemental Information). We then immediately obtain

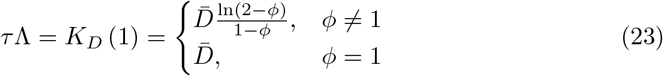

Since lim_*ϕ*→2_ *K*_*D*_ (1) =, 0 *< ϕ <* 2 is the range that the Fano factor of division counts can take within this scheme.

**Figure 3.**
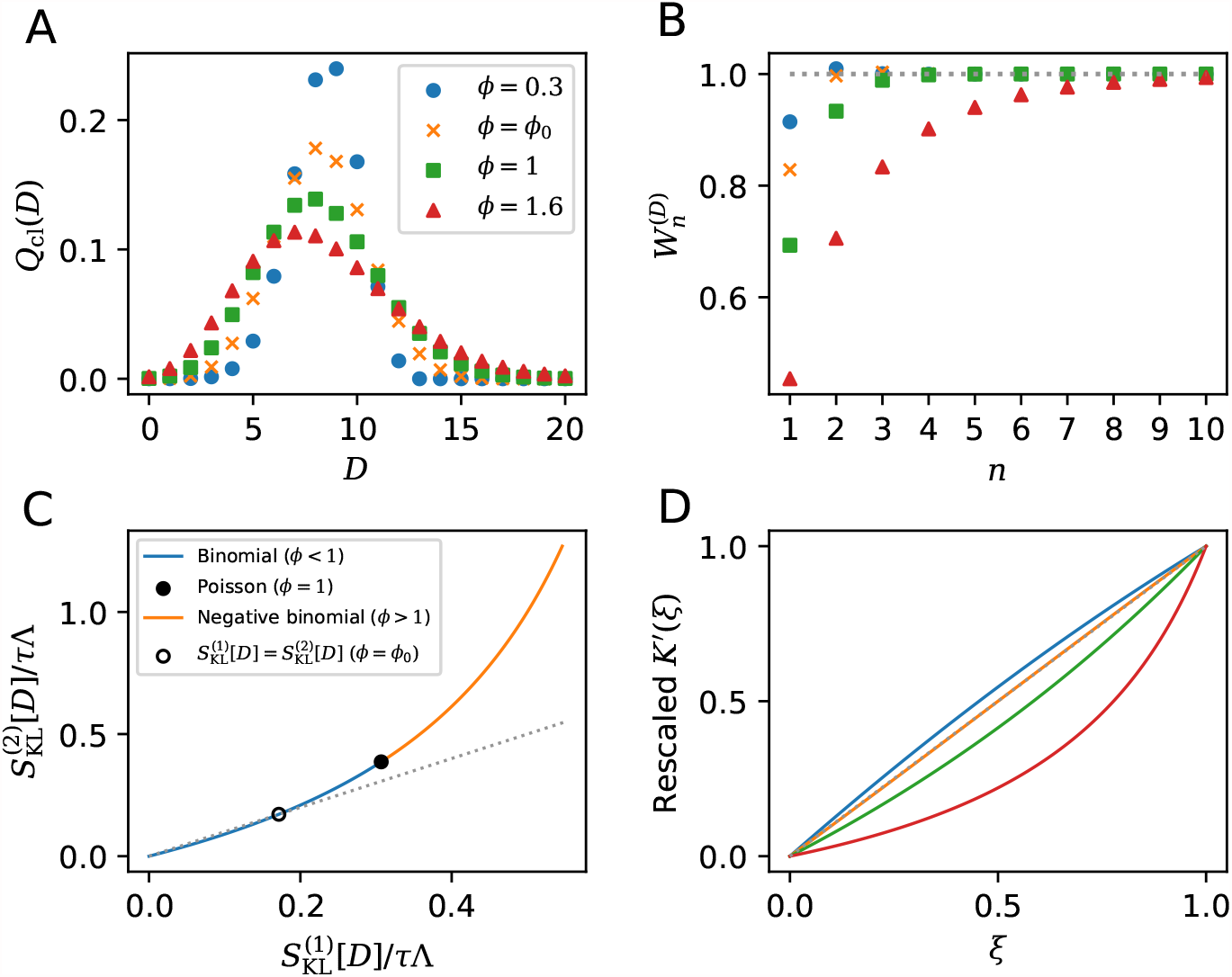
Analytical calculations of *K*_*D*_(*ξ*) and related relations given specific form of division count distributions. **A**. Chronological division count distributions. *ϕ* = 0.3 and *ϕ* = *ϕ*_0_(= 0.5857…) are binomial, *ϕ* = 1 is Poisson and *ϕ* = 1.6 is negative binomial. 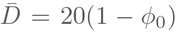 is fixed. **B**. Cumulative contributions of fitness cumulants. Parameter values are given in panel A legend. **C**. The relation between two selection strength measures. Binomial (blue curve), Poisson (closed black circle) and negative binomial (orange curve) are indicated on the single curve plotted using Eqs. 24 and 25 within the range of 0 *< ϕ <* 2. The point where 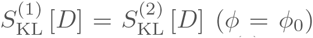 is indicated by the open black circle. The grey dotted line corresponds to 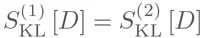 **D**. Convexity of 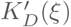. Y-axis shows a rescaling of 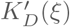 according to 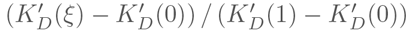. The same values of *ϕ* as in **A** are used; *ϕ* = 0.3 (blue), *ϕ* = *ϕ*_0_ (orange), *ϕ* = 1 (green) and *ϕ* = 1.6 (red). The grey dotted line indicates the case that 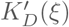 is a linear function of *ξ*.

Using (Eq. 22) allows us to calculate the cumulative contribution of fitness cumulants 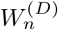 (Eq. 20). Plotting 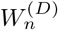 shows that the contribution of higher-order fitness cumulants becomes significant when *ϕ* is large (Fig. 3B). Also, evaluating the values of the derivative of Eq. 22 at *ξ* = 0 and *ξ* = 1, we have

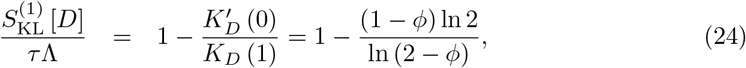

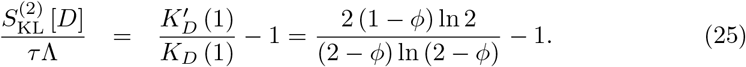

Therefore, 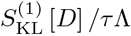 and 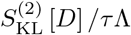 depend only on the Fano factor *ϕ*. In particular, 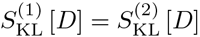 has 2 roots *ϕ* = 0, *ϕ*_0_(= 0.5857…); 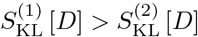 if 0*< ϕ < ϕ*_0_ and 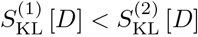 if *ϕ*_0_ *< ϕ <* 2 (Fig. 3C). Plotting 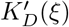 confirms that the covexity direction changes around *ϕ*_0_ (Fig. 3D). These analyses demonstrate how one can extract detailed information regarding selection in populations from *Q*_cl_ (*D*).

In Supplemental Information, we show the analytical calculation for a cellular population in which cells divide with gamma-distributed uncorrelated inter-division times. In this example, one can obtain *K*_*D*_(*ξ*) in the long-term limit condition, and the time-rescaled *n*-th order fitness cumulant is given as 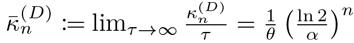, where *α* and *θ* are the shape and scale parameters of generation time distribution.

Therefore, unlike the central limit theorem, the contribution of higher-order fitness cumulants to population growth remains even in the long-term limit. Furthermore, this calculation demonstrates how the shape of the generation time distribution influences the cell population’s long-term growth rate.

### Relation of selection strength and population growth rate under fitness perturbations

We consider the response of population growth rate to perturbations that cause lineage fitness to change from *D*(*σ*) ln 2 to (1 − *ϵ*)*D*(*σ*) ln 2, and rewrite the population growth rate as

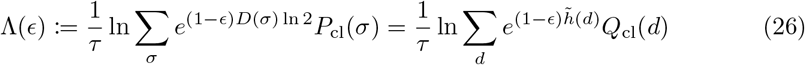

from (Eq. 2). We have Λ(0) = Λ, and note that 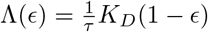 from (Eq. 15). Differentiating Λ(*ϵ*) with respect to *ϵ*, and evaluating at *ϵ* = 0, we find

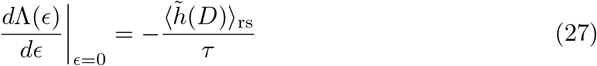

(see Supplemental Information). This relation shows that the change of population growth rate for small *ϵ* is proportional to the retrospective mean fitness of the unperturbed population. Since 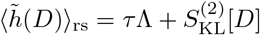 (Eq. 10), the relative change of population growth rate is

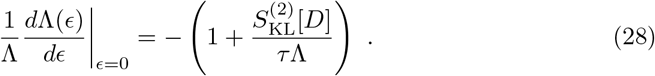

Therefore, a population with higher selection strength will exhibit a greater change in population growth rate upon perturbation. The selection strength measure 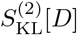 represents additional loss of population growth rate due to division count heterogeneity before perturbation since 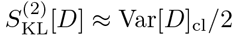 when the third or higher-order cumulants are negligible (see Supplemental Information).

It is, however, unclear what kind of perturbation *ϵ* represents and whether it has any realistic counterparts. As we see below, one manifestation of *ϵ* occurs via a cell removal operation. Consider the removal of a branch in the genealogical tree just after each cell division with the probability of 1 − 2^−*ϵ*^ (*ϵ >* 0) (Fig. 4A). In this case, the probability that a cell remains in the population after a cell division is 2^−*ϵ*^, and the growth of cell lineages that originally divided *d* times will be effectively reduced by the factor (2^−*ϵ*^)^*d*^. Consequently, the number of cell lineages that reach the end time point will also be effectively reduced from 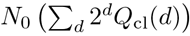 to 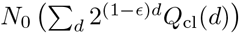. Therefore, the population growth rate under this branch removal operation is given by (Eq. 26), and the relative change of population growth rate is

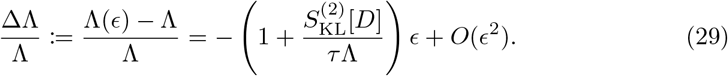

We validated this relation by simulating population growth with and without the cell removal operation (Fig. 4B-E and S1). We considered populations in which cells divide according to a gamma-distributed cell division time without mother-daughter correlations, with different values of the shape parameter which yield different selection strengths (Fig. 4B). The result confirmed that the relative changes of population growth rates by the probabilistic removal of cells followed 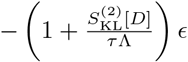 in all the conditions (Fig. 4C-E). We also tested this relation for cell populations with positive mother-daughter correlations of division intervals, which are often found for eukaryotic cells ([21–24]). We confirmed that the response relation was valid irrespectively of the strength of mother-daughter correlations (Fig. S1), which shows that the relation is general and independent of the specific dynamics of the cell division process. These results demonstrate that the response of population growth rate to cell the removal operation depends on the selection strength 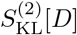 in the original population.

**Figure 4.**
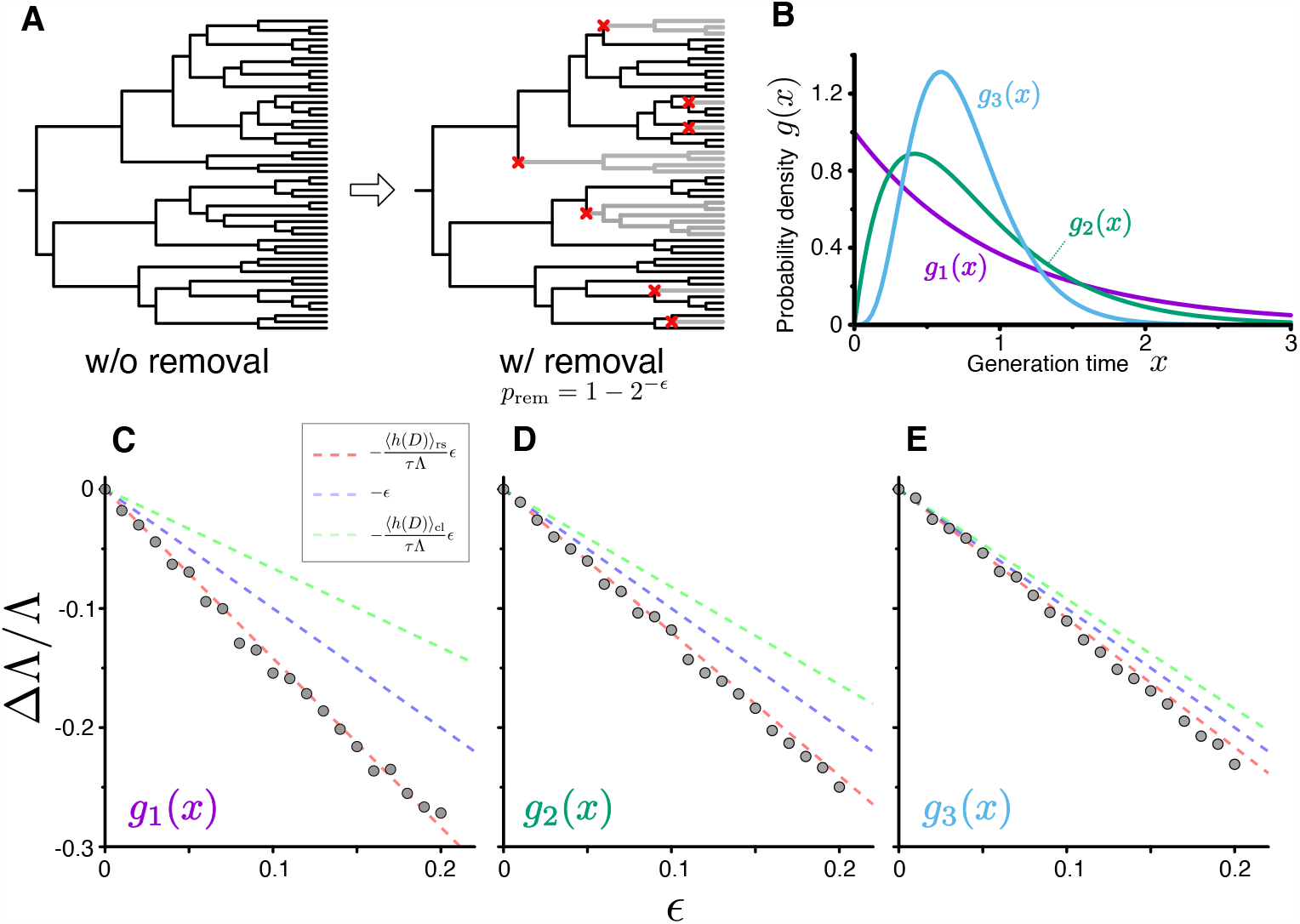
Population growth rate response to cell removal perturbation. **A**. Scheme of random cell removal. Here we consider the situation where cells were removed probabilistically after each cell division. Red crosses represent cell removal positions in the tree. The lineages after cell removal points disappear from the tree. Consequently, the number of cells at the end time point decreases. **B**. Generation time distributions used in the simulation. We assumed that cellular generation time follows gamma distributions in the simulation. We set the shape parameter to either 1 (*g*_1_(*x*)), 2 (*g*_2_(*x*)), or 5 (*g*_3_(*x*)). **C-E**. Population growth rate changes by cell removal perturbation. Gray points show the relative changes in population growth rate ΔΛ*/*Λ := (Λ(*ϵ*) − Λ(0))*/*Λ(0). Cell removal probability was set to 1 − 2^−ϵ^ in each condition of perturbation strength *ϵ*. Broken red lines represent the theoretical prediction 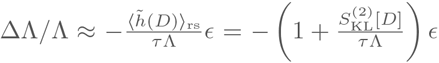. The lines of ΔΛ*/*Λ = −*ϵ* (blue) and 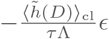 (green) are shown for reference. The generation time distributions used in the simulation are *g*_1_(*x*) for C, *g*_2_(*x*) for D, and *g*_3_(*x*) for E.

### Experimental evaluation of fitness cumulants’ contributions to population growth

Next, we apply this framework of cell lineage statistics to experimental single-cell lineage data of various organisms. The list includes bacterial cells (*Escherichia coli* and *Mycobacterium smegmatis*), unicellular eukaryotic cells (*Schizosaccharomyces pombe*), and mammalian cancer cells (L1210 mouse leukemia cells). As summarized in Table S1 and S2, we used cellular lineage data newly obtained in this study as well as other existing datasets [13, 22, 25, 26]. The *E. coli* and *S. pombe* data include several culture conditions to compare cumulants’ contributions to population growth across environments. The *E. coli* data were obtained using either agarose pad or the microchamber array microfluidic device, yielding genealogical tree information such as the one shown in Fig. 1. The *S. pombe* and L1210 cell data were obtained with mother machine microfluidic devices [6], which provide isolated cell lineage information but discard tree information due to its cell exclusion scheme. We assumed that these isolated cell lineages would follow chronological statistics and evaluated chronological distributions and selection strength according to the method described in Supplemental Information. All of the data analyzed in this section were taken from cell populations growing at approximately constant rates.

First, we evaluated the first-order cumulants’ contributions 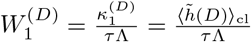 (Eq. 20), finding that 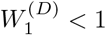 for all the samples and conditions (Fig. 5A). This result confirms that the chronological mean fitness cannot fully account for the population growth rates and that the contribution of division count heterogeneity must be taken into account. 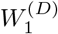 for *S. pombe* was consistently closer to 1 than those for the other cell types except one condition (EMM, 34°C), suggesting that *S. pombe*’s growth is less heterogeneous under most culture conditions.

**Figure 5.**
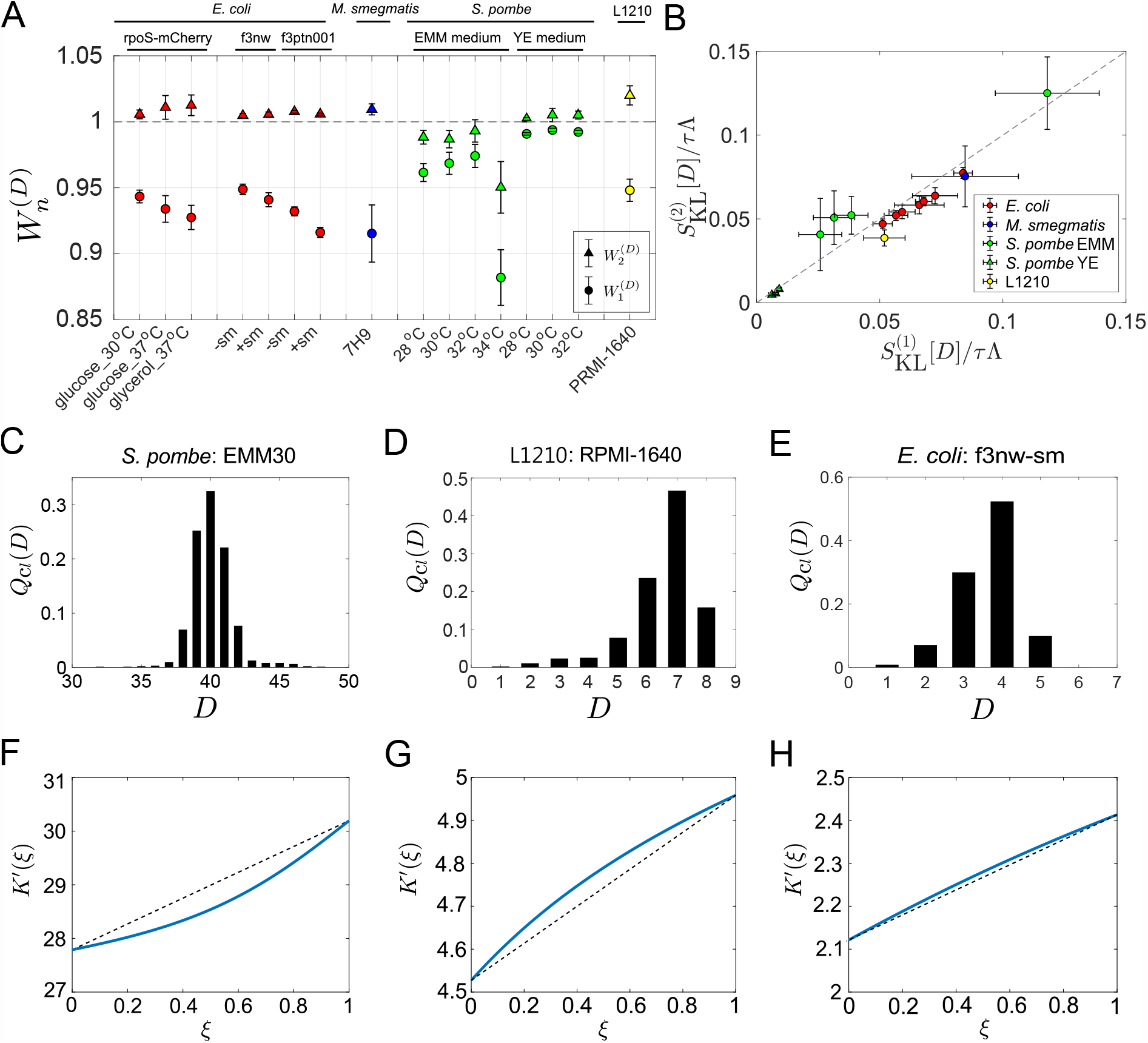
Application of cell lineage statistics to experimental data. **A**. Contributions of fitness cumulants to population growth. 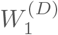 and 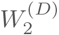 were evaluated for the experimental cell lineage data from *E. coli* (red), *M. smegmatis* (blue), *S. pombe* (green), and L1210 mouse leukemia cells (yellow). The *E. coli* rpoS-mcherry data were newly obtained in this study (see Materials and Methods). The other data were taken from literature: *E. coli* f3nw and f3ptn001 from [13]; *M. smegmatis* from [25]; *S. pombe* from [26]; and L1210 from [22]. Circles and triangles represent 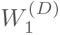 and 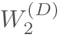, respectively. Error bars represent the two standard deviation ranges estimated by resampling the cellular lineages (see Materials and Methods). **B**. Relationship between 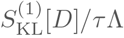 and 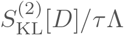. Colors correspond to the cellular species as in A. The *S. pombe* data were further categorized into two groups: Circles for the EMM conditions; and triangles for the YE conditions. **C-E**. Representative chronological distributions of division count, *Q*_cl_(*D*). **F-H**. Graphical representation of 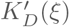. F for *S. pombe* EMM30; G for L1210 RMPI-1640; and H for *E. coli* f3nw-sm.

We next evaluated 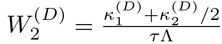, finding that 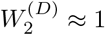 for most of the conditions (Fig. 5A). This result indicates small contributions of the third or higher-order cumulants to population growth. Consistent with this result, 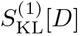 and 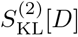 were almost identical in most conditions (Fig. 5B). Note that 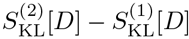 depends only on the third or higher-order cumulants (Eq. 21). The chronological distributions *Q*_cl_(*D*) of these samples were nearly symmetric in most cases, however, under the conditions where the deviations of 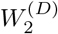 from 1 are larger, such as *S. pombe* in EMM medium and L1210, the distributions were skewed slightly (Fig. 5C-E, S2). Such distribution skew was reflected in the convexity directions of 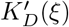-plots (Fig. 5F-H, S3). In *S. pombe* EMM medium conditions, 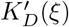 was convex downward in the interval 0 ≤ *ξ* ≤ 1 except for EMM 34°C (Fig. 5F and S3). Therefore, in contrast to a common assumption that selection necessarily decreases fitness variance, here we show that under certain conditions selection can increase fitness variance among cellular lineages.

### The contributions of higher-order cumulants become significant in the regrowth from a late stationary phase

We further applied the framework to the cell lineage data of *E. coli* populations regrowing from an early or late stationary phase. To conduct time-lapse observations of regrowing cell populations, we used a microfluidic device equipped with microchambers etched on a glass coverslip. We sampled *E. coli* cells either from an early or late stationary phase batch culture and enclosed the cells into the microchambers by a semipermeable membrane [10, 27]. We switched flowing media from stationary-phase conditioned medium to fresh medium at the start of time-lapse measurements and recorded the growth and division of individual cells (Fig. 6A, see Materials and Methods).

**Figure 6.**
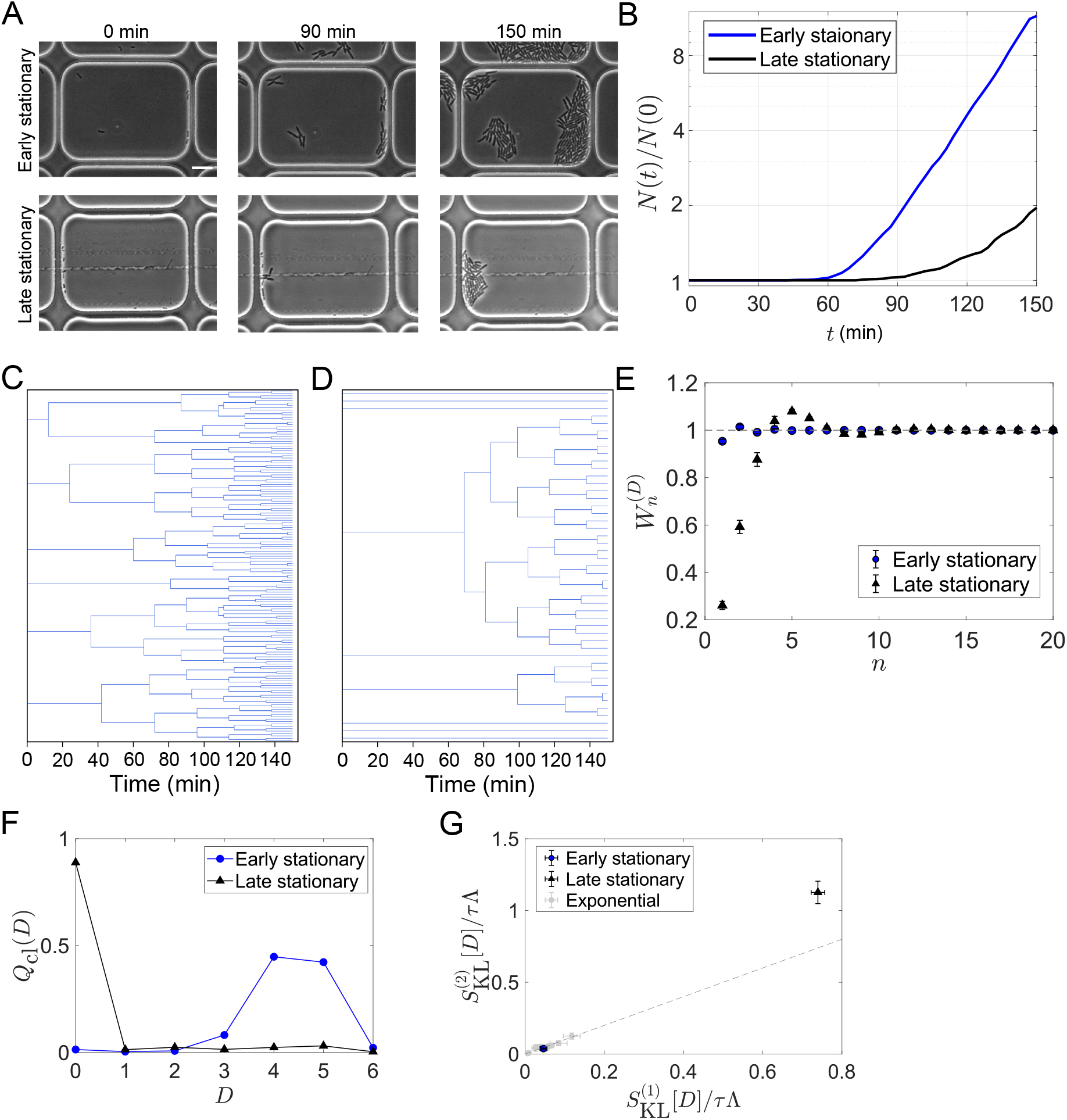
Strong selection in the *E. coli* population regrowing from a late stationary phase. **A**. Time-lapse images. Cellular regrowing dynamics from early and late stationary phases were observed by time-lapse microscopy. Cells were enclosed in the microchambers etched on coverslips. The top three images show representative images of the cells from an early stationary phase. The bottom three images show the cells from a late stationary phase. Scale bar, 5 *µ*m. **B**. Population dynamics. The number of cells at each time point normalized by the initial cell number (*N* (*t*)*/N* (0)) was plotted against time *t. N* (0) was 307 for the early stationary sample and 295 for the late stationary sample. **C, D**. Representative cellular lineage trees in the regrowing kineics from the early stationary phase (C) and the late stationary phase (D). The trees correspond to the time-lapse images in A. **E**. Cumulative contributions of fitness cumulants to population growth. Error bars represent the two standard deviation ranges estimated by resampling the cellular lineages (see Materials and Methods). **F**. Chronological distributions of division count *Q*_cl_(*D*). **G**. Relationships between 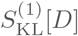 and 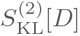. The blue and black points show the results for the early stationary phase sample and the late stationary phase sample, respectively. Gray points represent the results for the cell populations growing at approximately constant growth rates shown in Fig. 5B.

The growth curves reconstructed by counting the number of cells at each time point showed lags in regrowth (Fig. 6B). The lag time was shorter for the populations from the early stationary phase. The lineage tree structures in the cell populations were markedly different between the conditions (Fig. 6C and D). The tree structures were more uniform for the early stationary phase sample with multiple divisions in most cell lineages (Fig. 6C), whereas those for the late stationary phase sample were more heterogeneous, with 90% of cells showing no divisions within the observation time (Fig. 6D).

We analyzed these data and found 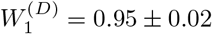 for the population from the early stationary phase and 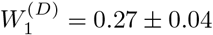 for the population from the late stationary phase (Fig. 6E). Therefore, the chronological mean fitness, 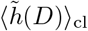, explains only 27% of the growth rate of the population regrowing from the late stationary phase. In other words, the selection strength contribution was 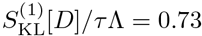. Thus, substantial selection existed in the cell population from late stationary phase. We also found that 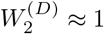 for the population from the early stationary phase, as observed for the *E. coli* populations growing at constant rates. In constrast, 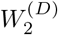 for the population from the late stationary phase was 0.61 ± 0.04, and 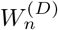 converged to 1 only after taking the cumulants up to approximately 10th-order into account (Fig. 6E). Reflecting the extreme skew to the right of the chronological distributions *Q*_cl_(*D*) (Fig. 6F), 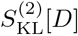 was significantly greater than 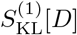 for the late stationary phase sample (Fig. 6G).

### Lineage statistics reveal condition-dependent fitness landscapes and selection strength for a growth-regulating sigma factor

RpoS is a sigma factor that controls the transcription of a large set of genes (10% of the genome) in *E. coli* [28]. High RpoS expression usually correlates with growth suppression; RpoS is induced when cells enter stationary phases or encounter stress conditions, such as starvation, low pH, oxidative stress, high temperature, or osmotic stress. Elevated RpoS expression provokes the intracellular programs to shut down growth and resist the stress [28]. However, it remains poorly understood how the continuum heterogeneity of RpoS expression levels is linked to the lineage fitness and selection in exponentially growing cellular populations. We therefore applied the lineage statistics framework to the single-cell time-lapse data of an *E. coli* strain expressing an RpoS-mCherry fusion protein from the native chromosomal locus and green fluorescent protein (GFP) from a low copy plasmid.

We quantified the time-scaled fitness landscapes *h*(*X*)*/τ* and relative selection strength *S*_rel_[*X*] (Eq. 14) under three growth conditions, taking the time-averaged mean fluorescent intensity of RpoS-mCherry or GFP along each cell lineage (proxies of time-averaged intracellular concentrations) as *X*. The result shows that the fitness landscapes and selection strength of RpoS expression level differ significantly among the growth conditions (Fig. 7). Under the glucose-37°C condition, the fitness landscapes of RpoS-mCherry and GFP expression were both decreasing functions (Fig. 7A and B). Thus, high expression of RpoS-expression and GFP in an exponentially growing population are both linked with lower lineage fitness. However, while the fitness landscape of GFP expression were nearly constant and showed significant decrease of fitness only at high expression levels, the fitness landscape of RpoS-mCherry decreased steadily in the observed expression range (Fig. 7A and B). Consequently, the relative selection strength for RpoS-mCherry was 2.6-fold larger than that for GFP (Fig. 7C).

**Figure 7.**
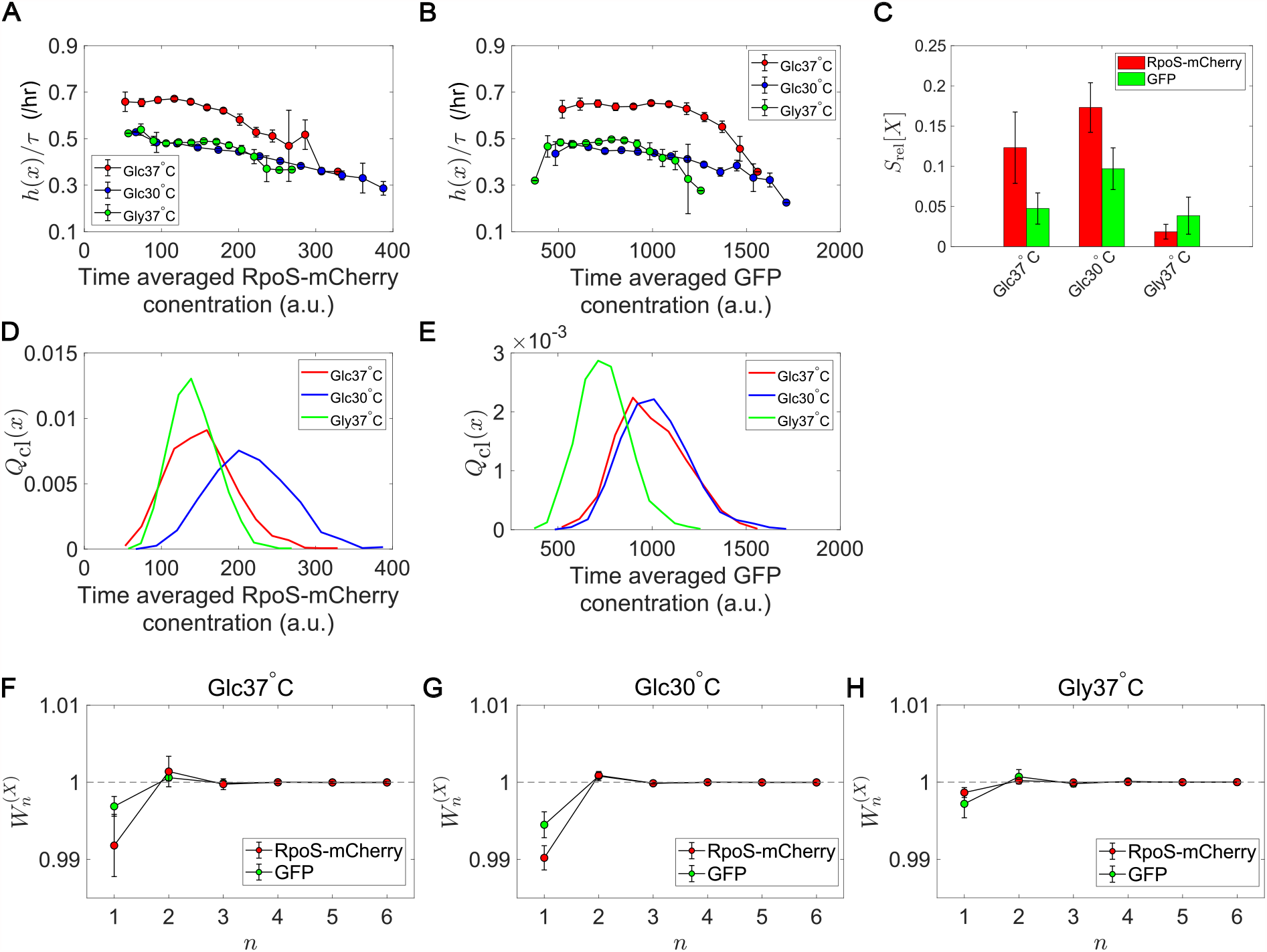
Fitness landscapes and selection strength for RpoS expression levels. A. Fitness landscapes for the time-averaged concentration (mean fluorescent intensity) for RpoS-mCherry. Fitness landscapes were scaled by the lineage length (observation duration) *τ*. Error bars represent the two standard deviation ranges estimated by resampling the cellular lineages. **B**. Fitness landscapes for the time-averaged concentration for GFP. **C**. Relative selection strength for the time-averaged concentrations of RpoS-mCherry (red) and GFP (green). **D, E**. Chronological distributions *Q*_cl_(*x*) for the time-averaged concentrations of RpoS-mCherry (D) and GFP (E). **F-H**. Cumulative contributions of fitness cumulants to population growth, 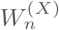, assuming that *X* is either time-averaged concentration of RpoS-mCherry (red) or time-averaged concentration of GFP (green). Error bars represent the two standard deviation ranges estimated by resampling the cellular lineages. Panel F is for the Glucose-37°C condition; Panel G for the Glucose-30°C condition; and Panel H for the Glycerol-37°C condition.

Under the glucose-30°C and glycerol-37°C conditions, the fitness landscapes for RpoS-mCherry level were also decreasing functions and close to each other but significantly downshifted from that for the glucose-37°C condition (Fig. 7A). This result reveals that cells could have different fitness for the same expression levels of RpoS, depending on the growth conditions. The selection strength for RpoS-mCherry was larger than that for GFP under the glucose-37°C and glucose-30°C conditions (Fig. 7C), which proves that the heterogeneity of RpoS expression in a population correlates with the lineage fitness more strongly than that of GFP under those conditions. On the other hand, the relative selection strength of RpoS-mCherry under the glycerol-37°C condition was the smallest among the three conditions and not significantly different from that of GFP (Fig. 7C). This is due to the relatively flat fitness landscapes in the central ranges of the distributions *Q*_cl_(*x*) (Fig. 7A and B) and the smaller variations of *x* in the population (Fig. 7D and E). These results reveal that the continuum heterogeneity of RpoS expression level in a population does correlate with the lineage fitness, but its contribution to selection depends on growth conditions.

We also evaluated the contributions of fitness cumulants for RpoS-mCherry expression to the population growth rate. Under all the conditions, 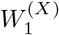 was lower than 1 (Fig. 7F-H). Therefore, the contributions of the higher-order fitness cumulants are non-negligible. However, the deviation of 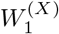 from 1 for RpoS-mCherry under the glycerol-37°C condition was small (Fig. 7H). Hence, in this growth condition, RpoS-mCherry expression barely correlated with fitness heterogeneity in the population.

## Discussion

Growth and division of individual cells are intrinsically variable, which causes division count heterogeneity among cellular lineages in a population. Such heterogeneity is ubiquitous across prokaryotic and eukaryotic cells, and its statistical properties could depend on the mechanisms and regulations determining cell division timings. Notably, division count heterogeneity influences population growth rate and, consequently, a population’s survival and evolutionary success. Therefore, understanding what statistical features are produced among cellular lineages and how these features contribute to population growth is essential for unraveling each organism’s survival and evolutionary strategy.

This report presents a cell lineage statistics framework to uncover the linkage between fitness distributions and population growth rate. We reveal that a population’s growth rate can be expanded by fitness cumulants of any lineage trait. The cumulant expansion allows us to quantify the contribution of each fitness cumulant, such as variance and skewness, to population growth rate. Applying this framework to the experimental cell lineage data revealed the fitness cumulants’ contributions to population growth for various organisms and environmental conditions. In particular, higher-order cumulants became significant in the regrowth of *E. coli* from a late stationary phase.

We remark that the cumulant expansion of population growth rate is valid only when all the cumulants are finite and when the Taylor expansion of *K*_*X*_ (*ξ*) around *ξ* = 0 also converges at *ξ* = 1. However, all the experimental data examined in this study exhibited stable convergence, including in the regrowth condition from the late stationary phase.

An advantage of this framework is its independence from any growth and division models. The mechanisms driving the growth and division of individual cells are diverse among organisms. For example, the properties of cellular growth and division, such as whether a cell’s size increases exponentially or linearly and whether cell size regulation follows sizer or adder models, could depend on cell types, organisms, and environmental conditions [29, 30]. Therefore, any model assumptions restrict applicability and necessitate model validation before application. The model independence of the framework presented here comes from the definitions of two essential quantities: the chronological and retrospective probabilities. Quantifying these probabilities requires only the information of the numbers of cells at initial and end time points and of division counts on each cellular lineage. Most importantly, division statistics is the focal information that connects molecular details underlying cellular growth and division to population growth. Regulatory mechanisms can influence population growth only by modulating the division statistics in a cellular population.

This framework is applicable even to cell populations growing under non-constant environmental conditions. We indeed utilized this framework to analyze the regrowth of growth-arrested cells from the stationary phase conditions. The selection strength contributions to population growth, 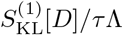, were below 10% in most cases under constant growth conditions. Nevertheless, it became over 70% in the regrowth of *E. coli* from the late stationary phase. While increased selection in non-constant environments may not be surprising itself, it is intriguing to ask how its contribution changes quantitatively depending on the conditions of environmental changes, such as nutrient upshift and downshift. The selection strength contribution in the regrowth from the early stationary phase was only 5%. This result clearly shows that how strongly selection acts in regrowing processes depends on stationary phase incubation durations. The framework is useful for unraveling and evaluating the structural differences of lineage trees quantitatively.

We identified the cellular populations in which selection acts to increase fitness variance in the retrospective statistics compared with the chronological statistics (Fig. 5F, 6G and S3). Such conditions are graphically recognized by the downward convexity of 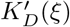 (Fig. 2). When the fourth or higher-order fitness cumulants are negligible, the convexity of 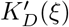 is determined primarily by the skewness of *Q*_cl_(*d*); positive skew of *Q*_cl_(*d*) with a long right tail makes 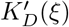 convex downward and 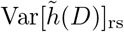 greater than 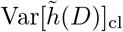. This consequence is intuitively understandable since the right tail of *Q*_cl_(*d*) is accentuated in proportion to *e*^*D*^ by selection, which leads to greater variance of *Q*_rs_(*d*). On the other hand, when the skew is negative with the long left tail, the effect of applying *e*^*D*^ is to diminish the tail and compress the distribution toward the fittest lineages.

We showed that division count heterogeneity among cellular lineages has dual facets: increasing population growth rate while sensitizing populations to perturbations. These two effects are quantitatively represented by 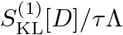 and 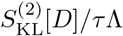, respectively. Therefore, the difference between these selection strength measures gauges which aspect of growth heterogeneity is more significant in the population.

Even though 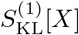 and 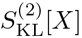 are different in general, the analysis revealed that they were nearly identical in most of the cellular populations growing at constant rates (Fig. 5). This result might suggest that, from a practical viewpoint, the contribution of higher-order cumulants becomes negligible under steady growth conditions, and the significant difference between 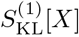 and 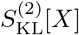 could be used as a probe for the non-stationarity of the population growth. This speculation must be examined experimentally using various organisms and cell types across diverse environmental conditions.

This framework is premised on complete lineage tree information. However, many methods of single-cell measurements continuously exclude cells from observation areas and provide only a part of the tree information. Therefore, extending this framework so that one can infer both chronological and retrospective probabilities from incomplete tree information is an essential future research direction. In this study, we calculated the fitness landscapes and selection strength measures for the cell lineage data obtained with the mother machine devices, assuming that these cell lineages would follow the chronological statistics. Such a simple approach is not yet available for larger-scale lineage tree data obtainable with the other single-cell measurement devices such as dynamics cytometer [10] and chemoflux [31]. Furthermore, it has been shown that the inference precision of population growth rate has non-monotonic dependence on the length of cell lineages obtained with mother machine devices [15]. Even though the difficulties to overcome are present, a comprehensive framework may permit a unified treatment of cellular lineage data obtained using various single-cell measurement methods.

Phenotypic heterogeneity is widely observed in diverse cellular systems, including both prokaryotic and eukaryotic cells. It is often considered that phenotypic heterogeneity allows bet-hedging against unpredictable environments and promotes the survival of cellular population [32]. However, quantitative evaluation of correlations between the traits of interest and fitness is usually an intricate problem. The cell lineage statistical framework described in this study offers a straightforward procedure applicable to any cellular genealogical data, which are now becoming increasingly available for various biological phenomena, including cancer metastasis [33] and stem cell differentiation [34–36]. Another important advantage of this framework is that it allows decomposing a population growth rate into chronological fitness and selection strength. It is thus intriguing to apply this framework to long-term evolutionary dynamics and quantify how the contributions of chronological mean fitness and selection evolve underlie the transitions of population growth rate. Such analysis might clarify the crucial roles of phenotypic heterogeneity in facilitating evolution.

## Materials and Methods

### Microfabrication of microchamber array

We constructed and used a microchamber array for conducting single-cell time-lapse observation under controlled environmental conditions. A microchamber is a well etched on a glass coverslip. We used two types of microchamber array. One is an array of microchamber, whose dimension is 70 *µ*m (w) × 55 *µ*m (h) × 1 *µ*m (d). This microchamber has a 21-*µ*m × 7-*µ*m pillar for supporting the membrane in the middle. We used this microchamber array for the exponential-phase experiment of *E. coli*. Another is an array of microchamber, whose dimension is 40 *µ*m (w) × 30 *µ*m (h) × 1 *µ*m (d). We used this type of microchamber array for the stationary-phase-regrowth experiment in Fig. 6. We fabricated these microchamber arrays following similar procedures described in [10, 27].

The photomasks for the microchamber array were created by laser drawing (DDB-201-TW, Neoark) on mask blanks (CBL4006Du-AZP, CLEAN SURFACE TECHNOLOGY). The photoresist on mask blanks was developed in NMD-3 (Tokyo Ohka Kogyo). The uncovered chromium (Cr)-layer was removed in MPM-E30 (DNP Fine Chemicals), and the remaining photoresist was removed by acetone. Lastly, the slide was rinsed in MilliQ water and air-dried.

The microchamber array was created in glass coverslips by chemical etching. First, we coated a 1,000-angstrom Cr-layer on a clean coverslip (NEO Micro glass, No. 1., 24 mm × 60 mm, Matsunami) by evaporative deposition and AZP1350 (AZ Electronic Materials) by spin-coating on the Cr-layer. We transferred the photomask patterns using a mask aligner (MA-20, Mikasa). After developing the photoresist in NMD-3 and the Cr-layer in MPM-E30, the coverslip was soaked in buffered hydrofluoric acid solution (110-BHF, Morita Kagaku Kogyo) for 14 minutes 20 seconds at 23°C for glass etching. The etching reaction was stopped by soaking the coverslip in milliQ water. The remaining photoresist and the Cr-layer were removed by acetone and MPM-E30, respectively.

### Fabrication of PDMS pad

We used a polydimethylsiloxane (PDMS) pad to flow culture medium and control the environmental conditions around the cells in the microchamber array. The PDMS pad was designed to have a square bubble-trap groove, which prevents interference with bright-field microscopic imaging by air bubbles in flowing media.

To create a mold for the bubble-trap groove, we spin-coated SU-8 3050 (Kayaku Advanced Materials) on a silicon wafer (ID 447, *ϕ* = 76.2 mm, University Wafer) and baked it at 95°C for 2 hours on a hot plate. The SU-8 layer was exposed to UV light on a mask aligner using a photomask and postbaked at 95°C for 2 hours. After cooled down to room temperature, the SU-8 photoresist was developed in the SU-8 developer (Kayaku Advanced Materials) and rinsed with isopropanol (Wako).

Part A and Part B of PDMS resin (SYLGARD 184 Silicone Elastomer Kit, DOW SILICONES) were mixed at 10:1 and poured onto the SU-8 mold. The air bubbles were removed under a decreased pressure for 30 min. The PDMS was cured at 65°C for 1 hour, and 20 mm × 20 mm square PDMS pad was cut out using a blade. We punched out two holes (*ϕ* = 2 mm) in the PDMS pad for the inlet and outlet, and 10-cm silicone tubes (SR-1554, Tigers Polymer Corp., outer *ϕ* = 2 mm, inner *ϕ* = 1 mm) were inserted into the holes. The tubes were fixed to the holes by gluing a small amount of PDMS around the tubes at the holes. This PDMS pad was washed in isopropanol by sonication and autoclaved for the single-cell measurements.

### Chemical decoration of coverslip and cellulose membrane

We washed the microfabricated coverslips by sonication in contaminon (Wako), ethanol (Wako), and 0.1 M NaOH solution (Wako). The washed coverslips were rinsed in milliQ water by sonication and dried at 140°C for 30 min. The washed coverslip was soaked in 1% (v/v) 3-(2-aminoethylaminopropyl)trimethoxysilane solution (Shinetsu Kagaku Kogyo) for 30 min and incubated at 140°C for 30 min to create an amino group on the glass surface. The treated coverslip was washed in milliQ water for 15 min and dried at 140°C for 30 min. 1 mg NHS-LC-LC-Biotin (Funakoshi) was dissolved in 25 *µ*l dimethyl sulfoxide and dispersed in 1 ml phosphate buffer (0.1 mM, pH8.0). 200 *µ*l of this biotin solution was placed on the coverslip and incubated at room temperature for four hours. The biotin solution was removed by soaking the coverslip in milliQ water.

We prepared a streptavidin-decorated cellulose membrane to enclose cells in the microchamber array while retaining a flexible environmental control. First, a 3 cm × 3 cm square cellulose membrane (Spectra/Por7 Pre-treated RC Tubing MWCO:25kD) was cut out and washed in milliQ water for 10 min. The membrane was incubated in a 0.1 M NaIO_4_ solution with gentle shaking for 4 hours at 25°C. After the wash in milliQ water, the treated membrane was incubated in a 500-*µ*l solution of streptavidin hydrazide (Funakoshi) (10 *µ*g/ml, dissolved in 0.1 mM phosphate buffer (pH7.0)) with gentle shaking for 14 hours at 25°C. The membrane was again washed in milliQ water and stored at 4°C.

### *E. coli* strains

We used two *E. coli* strains: MG1655 and MF3 *rpoS-mcherry* (MG1655 Δ*fliC* Δ*fimA* Δ*flu rpoS-mcherry* /pUA66-P_rplS_-*gfp*). MG1655 was used in the regrowth experiment from the stationary phases (Fig. 6). MF3 *rpoS-mcherry* was used for analyzing the growth in constant environments (Fig. 5 and 7). In MF3 *rpoS-mcherry*, the three genes, *fliC, fimA*, and *flu*, were deleted, and *mcherry* gene was inserted downstream of *rpoS* gene to express RpoS-mCherry translational fusion protein. This strain also expresses green fluorescent protein (GFP) from a low-copy plasmid, pUA66-P_rplS_-*gfp*, taken from a comprehensive library of fluorescent transcriptional reporters [37].

### Culture conditions and sample preparation (exponential growth)

We used MF3 *rpoS-mcherry E. coli* strain and cultured the cells in M9 minimal medium (Difco) supplemented with 1/2 MEM amino acids solution (SIGMA) and 0.2% (w/v) glucose or glycerol as a carbon source. We set the cultivation temperature either at 37°C or 30°C.

To prepare *E. coli* cells for single-cell observation, we first inoculated a glycerol stock into a 3 ml culture medium and incubated it with shaking overnight under the same conditions of culture medium and temperature as those used in the time-lapse measurement. 30 *µ*l of the overnight culture was inoculated in a 3 ml fresh medium and incubated with shaking until the optical density at *λ* = 600 nm reaches 0.1-0.3. This exponential-phase culture was diluted to OD_600_ = 0.05, and 0.5 *µ*l of the diluted cell suspension was spotted on the microchamber array on a biotin-decorated coverslip. A 5-mm × 5-mm streptavidin-decorated cellulose membrane was placed gently on the cell suspension on the coverslip, and an excess cell suspension was removed by a clean filter paper. A small piece of agar pad made with the culture medium and 1.5% (w/v) agar was placed on the cellulose membrane to maintain the culture conditions around the cells until tight streptavidin-biotin bonding was formed between the coverslip and the membrane. After 5-min incubation, the agar pad was removed, and the PDMS pad for medium perfusion was attached on the coverslip via a square-frame two-sided seal (Frame-Seal™ Incubation Chambers, Bio-rad). We immediately filled the device with the fresh medium and connected it to a syringe pump on the microscope stage.

### Culture conditions and sample preparation (regrowth from stationary phases)

We used *E. coli* MG1655 strain and cultured the cells in Luria-Bertani (LB) medium at 37°C. To prepare the cells for the time-lapse experiment, a glycerol stock of this strain was inoculated into a 2 ml LB medium and cultured with shaking for 15 hours. The cell culture was diluted in 50 ml fresh LB medium to OD_600_ = 0.005 and again cultured with shaking as a pre-culture. For preparing the early-stationary-phase conditioned medium, 7 ml pre-culture cell suspension at 8 hours (OD_600_ *≈*4.3) was spun down at 2,600 G for 12 min. The supernatant was filtered through a 0.22-*µ*m filter. For preparing cells for time-lapse observation, a 10-*µ*l pre-culture cell suspension at 8 hours was mixed with 240 *µ*l early-stationary-phase conditioned medium. 1-*µ*l of this diluted cell suspension was placed on the microchamber array on a biotin-decorated glass coverslip. A 5-mm × 5-mm streptavidin-decorated cellulose membrane was placed gently on the cell suspension on the coverslip, and an excess cell suspension was removed by a clean filter paper. A small piece of a conditioned medium agar pad made with 1.5% (w/v) agar was placed on a cellulose membrane to maintain the early stationary phase condition during the incubation. After 5-min incubation, the conditioned medium agar pad was removed, and the PDMS pad for medium perfusion was attached on the coverslip via a square-frame two-sided seal. We immediately filled the device with the conditioned medium and connected it to a syringe pump. We maintained the chamber filled with the conditioned medium until we started the time-lapse observation. The conditioned medium was flushed away immediately before starting the time-lapse measurement by flowing fresh LB medium. After flowing 2 ml fresh LB medium at 32 ml/h, the flow rate was decreased and maintained at 2 ml/h throughout the time-lapse measurement.

We followed the same procedures for the late stationary phase sample except that we sampled the cells and prepared the conditioned medium from a 24-hour pre-culture cell suspension (OD_600_ *≈* 3.0).

### Time-lapse measurements and image analysis

We used Nikon Ti-E inverted microscope equipped with Plan Apo *λ* × 100 phase contrast objective (NA1.45), ORCA-R2 cooled CCD camera (Hamamatsu Photonics), Thermobox chamber (Tokai hit, TIZHB), and LED excitation light source (Thorlabs, DC2100). The microscope was controlled by Micromanager [38]. In the exponential phase experiments, we monitored 25-30 microchambers in parallel in one measurement and acquired the phase-contrast, RpoS-mCherry fluorescence, and GFP fluorescence images from each position with a 3-min interval. We repeated the time-lapse measurement for each culture condition three times. In the regrowth experiment from the stationary phases, we monitored 150-250 microchambers in parallel with a 3-min interval and acquired only phase-contrast images.

We analyzed the time-lapse images by ImageJ [39]. We extracted the information of cell size (projected cell area), RpoS-mCherry fluorescence mean intensity, and GFP fluorescence mean intensity of individual cells along with division timings on each cell lineage for the exponential phase experiment. We extracted only division timings on each cellular lineage for the regrowth experiments from the stationary phases and used this information for further analysis.

## Data analysis

### Distributions and selection strength measures for division count

We calculated the distributions and selection strength measures of {*D*} as follows. With the list of division counts *D* for each lineage *σ*, the chronological and retrospective probabilities were evaluated as *P*_cl_(*σ*) = 2^−*D*(*σ*)^*/N*_0_ and *P*_rs_(*σ*) = 1*/N*_*τ*_, respectively, where *N*_0_ is the number of cells at *t* = 0 and *N*_*τ*_ is that at *t* = *τ*. From these probabilities, the chronological and retrospective distributions of *D* were obtained by summing the lineage probabilities for each division count, i.e.,

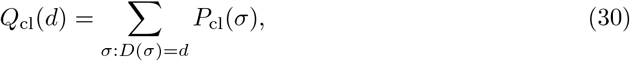

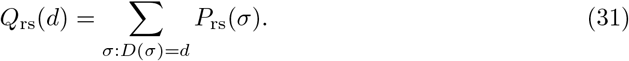

The selection strength measures, 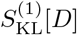 and 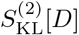, were calculated as

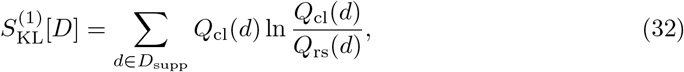

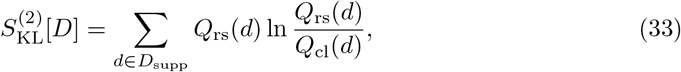

where *D*_supp_ is the support of both chronological and retrospective probabilities with respect to *D*, which is common between the two probabilities.

### Distributions and selection strength measures for time-averaged fluorescence intensity of RpoS-mCherry and GFP

We obtained the mean fluorescence intensity of RpoS-mCherry and GFP along with the genealogical trees in the time-lapse measurements of *E. coli* MF3 *rpoS-mcherry* strain. We analyzed the time-averaged fluorescence intensity of RpoS-mCherry and GFP as a lineage trait *X* and evaluated their distributions, fitness landscapes, and selection strength measures (Fig. 7). For each cell lineage, the time-averaged fluorescence intensity was calculated as

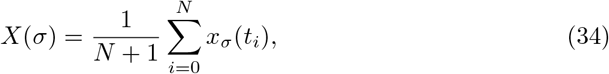

where *t*_*i*_ = *t*_start_ + *i*Δ*t* min (*t*_start_ is the start time of the cell lineage; Δ*t* = 3 min is the time-lapse interval), and *x*_*σ*_(*t*_*i*_) is the mean fluorescence intensity at time *t*_*i*_.

To calculate the chronological and retrospective probability densities and the fitness landscapes, we set the bin width *ΔX* to

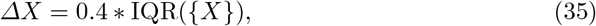

where IQR({*X*}) is the interquartile range of the set of *X*(*σ*) from all the cell lineages. Then, the interval was defined as 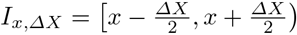 for *x* = min({*X*}), min({*X*}) + *ΔX, …*, min({*X*}) + (*L* − 1)*ΔX*, where *L* is the number of total bins given by 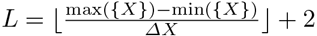.

We calculated the chronological and retrospective probability distributions of *X* by

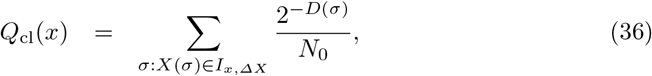

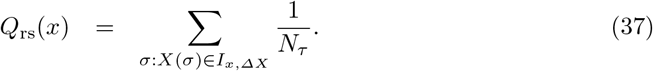

The fitness landscape *h*(*x*) was evaluated by

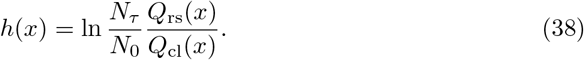

The selection strength measures were evaluated by

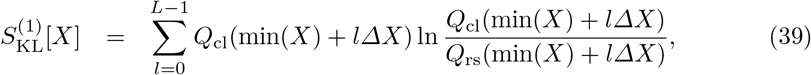

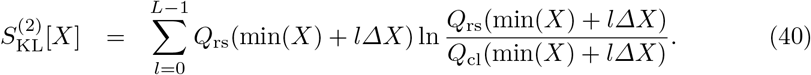

### Cumulant generating functions and cumulants

To plot the differential of the cumulant generating functions in Fig. 5F-H, we evaluated 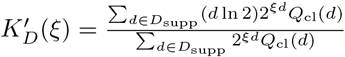 by changing *ξ* from 0 to 1 with the step size 0.01.

We calculated the cumulative contributions of fitness cumulants to the population growth 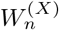 (Fig. 5A, 6E, and 7F-H) using a julia package, JuliaDiff/TaylorSeries.jl (https://github.com/JuliaDiff/TaylorSeries.jl).

### Error estimations by resampling method

To evaluate the error ranges of the quantities calculated in the analysis, we created 20000 randomly resampled datasets for each condition and reported the means and two standard deviation ranges in the results.

For the datasets of colony growth (*E. coli* and *M. smegmatis*), *N*_*τ*_ lineages were randomly sampled with replacement according to the probability weight *P*_rs_(*σ*) for each resampled dataset. In each resampled dataset, the initial number of cells was estimated as 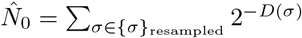

For the datasets taken using the mother machines (*S. pombe* and L1210), we randomly sampled *N*_0_ lineages with an equal weight, which corresponds to the chronological probability in this setting. *N*_*τ*_ was estimated as 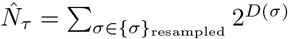

### Simulating the effect of cell removal on population growth rates

We simulated cell population growth with cell removal using a custom C script. The gamma distributions were adopted as generation time distributions. We assigned the shape parameter to *k* =1, 2, or 5 and the scale parameter to *θ* = 2^1*/k*^ −1. The perturbation strength *E* was changed from 0 to 0.2 with the interval 0.01.

As a pre-run, we started a simulation from a newborn cell and assigned its generation time randomly according to a pre-defined gamma probability distribution. We assumed that this cell divided into two daughter cells at the end of the generation. Each daughter cell was removed with probability 1− 2^*−ϵ*^ and assigned with generation time from the same pre-defined probability distribution if it escaped removal. Repeating this procedure, we let the population grow until all of the remaining cell lineages in the population exceed the maximum duration *T*_max_ = 8.0. The time to the next division of each cell lineage at *T*_max_ was exported as the first division time in the main simulation. This pre-run was repeated 1,000 cycles to export a sufficiently sizable list of first division times.

In the main simulation, we started from a progenitor cell with its division time randomly assigned from the first division time list exported in the pre-rum. For the daughter cells born from the first divisions and their descendants, the assignment of generation time and the cell removal were done as in the pre-run. We stopped further production of daughter cells in each lineage if it exceeded *T*_max_ = 8.0. We repeated this main simulation 1,000 cycles starting from different progenitor cells. The number of cell divisions in each cell lineage until *T*_max_ was exported for analysis.

We calculated the population growth rate at each perturbation strength as

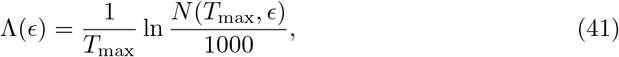

where *N* (*T*_max_, *ϵ*) is the number of cell lineages at *T*_max_ when the perturbation strength was *ϵ*. The chronological and retrospective mean fitness of division count without cell removal was calculated as

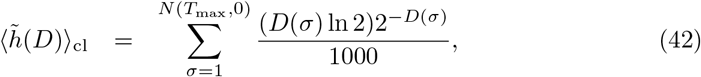

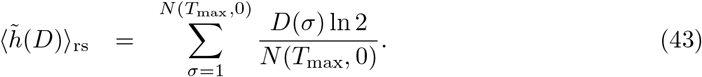

When simulating the cell population with mother-daughter correlation time, we randomly assigned the generation time from the gamma probability distribution with its shape parameter 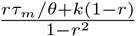 and scale parameter (1− *r*^2^)*θ*, where *τ*_*m*_ is the generation time of the mother cell, *r* is the correlation coefficient of generation time between neighboring generations. The stationary distribution of this transition probability approximates the gamma distribution with shape parameter *k* and scale parameter *θ* to good precision with identical first and second-order moments irrespective of the parameters *k, θ*, and *r*. In Fig. S1, we fixed *k* = 2 and 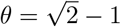 and set *r* to 0, 0.2, 0.4, or 0.6.

## Acknowledgments

We thank Tetsuya J Kobayashi and the members of the Wakamoto Lab for discussion. This work was supported by JST CREST Grant Number JPMJCR1927 (Y.W.); JST ERATO Grant Number JPMJER1902 (Y.W.); NIH Grant Number R01-GM097356 (E.K.); and Japan Society for the Promotion of Science KAKENHI Grant Number 17H06389 and 19H03216 (Y.W.).

## Supplemental Information

### Supplemental text

#### The properties of the selection strength of division count

Below we derive several important properties of the selection strength of division count. We focus on the selection strength measure 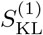 and write it as *S* this section for conciseness. However, the conclusions are likewise valid for *S*_JF_ and 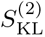.

The most detailed description of cellular lineage statistics is based on individual lineages *σ*. We define the selection strength of cellular lineages as

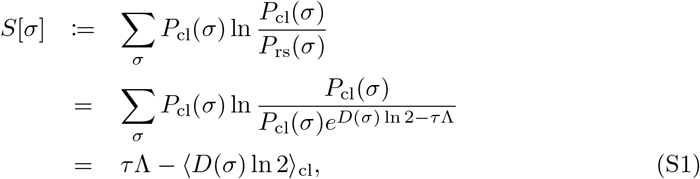

where ⟨*D*(*σ*) ln 2⟩_cl_ = Σ_*σ*_ (*D*(*σ*) ln 2)*P*_cl_(*σ*).

From the definition of fitness landscape (Eq. 3),

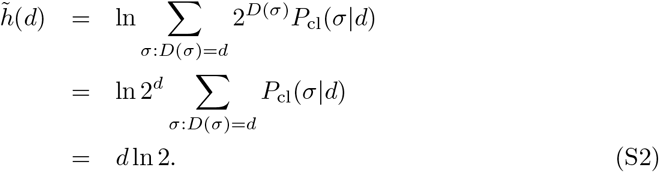

On the other hand,

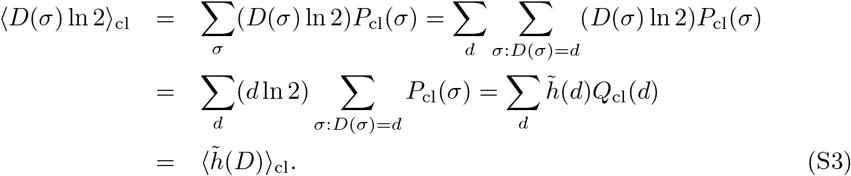

This proves that the chronological mean fitness of cellular lineages equals the chronological mean fitness of division count.

Since 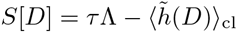 and *S*[*σ*] = *τ* Λ − *(D*(*σ*) ln 2)_cl_ (Eq. 11 and S1),

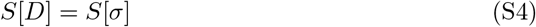

is also held. This result shows that the selection strength of *D* is equivalent to the selection strength of cellular lineages despite *D* being a coarse-grained lineage trait.

Another important property of *S*[*D*] is that it sets the maximum bound for the selection strength of any lineage traits. Now we consider the joint probability distributions of *D* and lineage trait *X*, which we write *Q*_cl_(*d, x*) and *Q*_rs_(*d, x*). We define the joint selection strength as

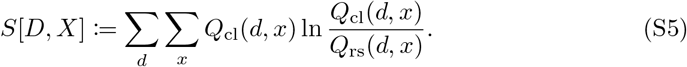

Using *Q*_cl_(*d, x*) = *Q*_cl_(*d*|*x*)*Q*_cl_(*x*) and *Q*_rs_(*d, x*) = *Q*_rs_(*d*|*x*)*Q*_rs_(*x*),

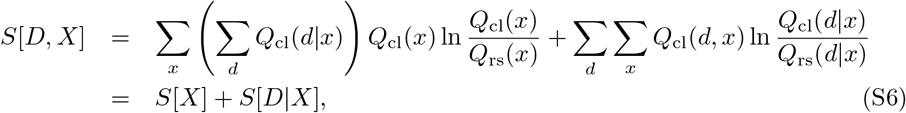

where 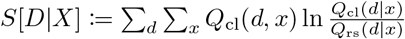, and we used ∑_*d*_ *Q*_cl_(*d*|*x*) = 1. Likewise, *S*[*D, X*] can also be decomposed as

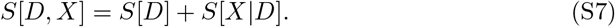

However, *S*[*X*|*D*] = 0 because

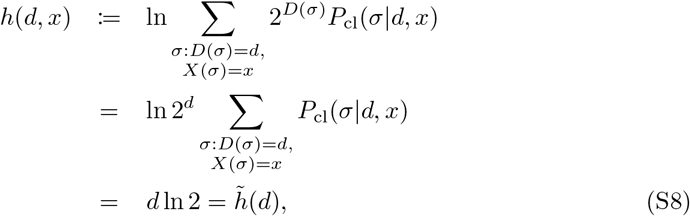

and

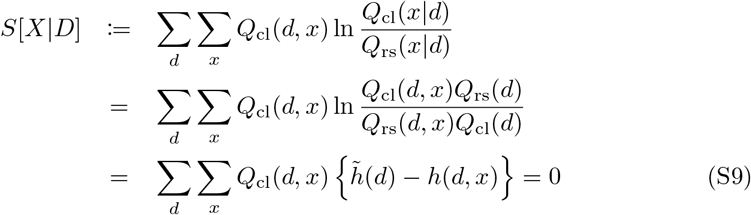

from (Eq. 4) and (Eq. S8). This leads to

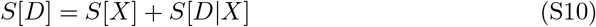

from (Eq. S6) and (Eq. S7). Furthermore, *S*[*D*|*X*] ≥ 0 from Jensen’s inequality. Thus,

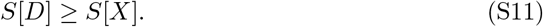

The equality is held when *D* is a deterministic function of *X*. This inequality shows that *S*[*D*] (= *S*[*σ*]) sets the maximum bound for the selection strength of any lineage trait *X*.

#### The cumulant generating function *K*_*X*_ (*ξ*) provides both chronological and retrospective fitness cumulants

In the main text, we introduced the cumulant generating function of *h*(*x*) with respect to the chronological distribution *Q*_cl_(*x*),

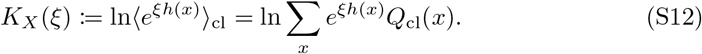

This function can also be written as

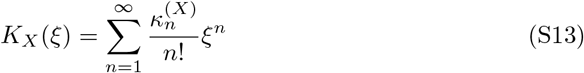

when the fitness cumulants 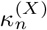 are all finite, and the Taylor expansion converges at *ξ*. Also,

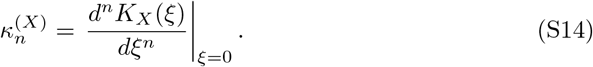

Below we prove that *K*_*X*_ (*ξ*) also gives the fitness cumulants on the retrospective distributions.

We define a cumulant generating function on the retrospective probability as

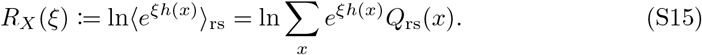

This function can be expanded by the fitness cumulants of the retrospective statistics 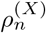 as

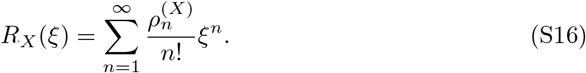

Therefore,

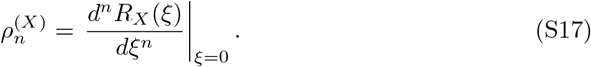

For example, 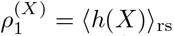 and 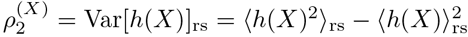.

Inserting *Q*_rs_(*x*) = *e*^*h*(*x*)−*τ*Λ^*Q*_cl_(*x*) into (Eq. S15),

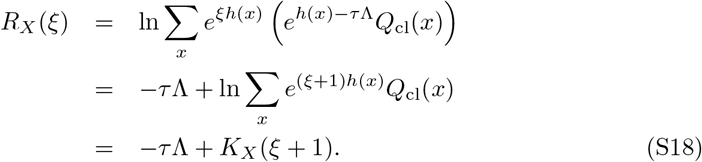

Hence,

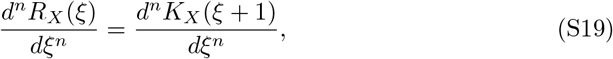

for *n* ≥1. This relation proves that evaluating 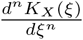 at *ξ* = 1 gives the *n*-th order fitness cumulant on the retrospective statistics; i.e.,

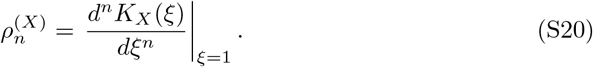

Furthermore, this leads to

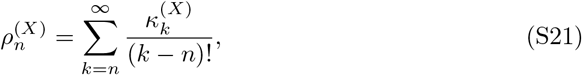

from (Eq. S13) and (Eq. S20). Similarly, evaluating (Eq. S19) at *ξ* = −1 gives

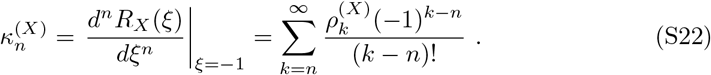

Analogously to (Eq. 18), we can also expand the population growth rate in terms of the retrospective cumulants, by evaluating (Eq. S18) at *ξ* = −1,

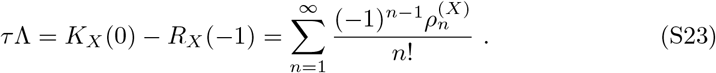

For example, when the fitness distribution is Gaussian for the chronological statistics,

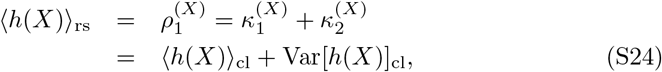

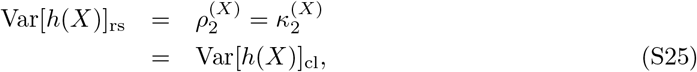

since 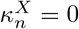 for *∀n* ≥ 3.

These results confirm that the function *K*_*X*_ (*ξ*) contains the information of both chronological and retrospective statistics.

#### Relationships between fitness cumulants and selection strength measures

In the main text, we have shown that the selection strength 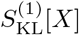 corresponds to the contribution of the second or higher-order fitness cumulants to population growth, i.e.,

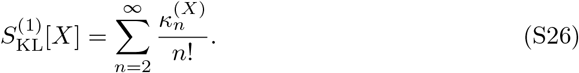

or alternatively, by substituting (Eq. S22) and (Eq. S23) we obtain

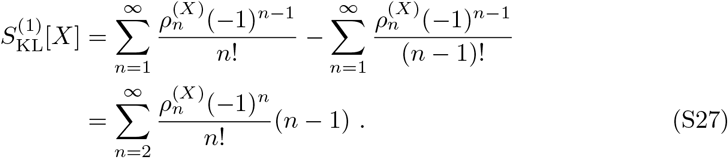

Similar expressions can also be found for 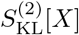. Since 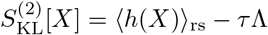 (Eq. 12), substituting (Eq. S23) yields

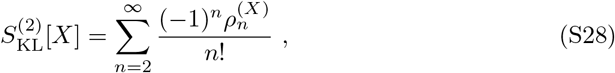

or alternatively, by substituting (Eq. S21) and (Eq. 18) we obtain

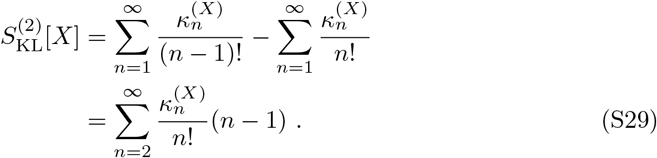

These show that both of 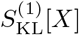 and 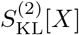 can be expanded by the chronological or retrospective fitness cumulants.

The difference between these two selection strength measures is

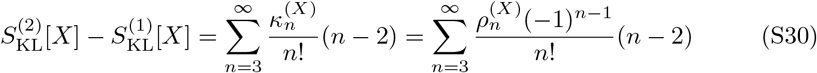

from (Eq. S26) to (Eq. S29). Thus, it depends only on the third or higher-order fitness cumulants.

Finally, another selection strength measure *S*_JF_[*X*] can also be expanded by the fitness cumulants as

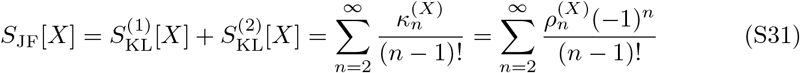

from (Eq. S26) to (Eq. S29). When the chronological fitness distribution is Gaussian (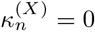 for *∀n* ≥ 3),

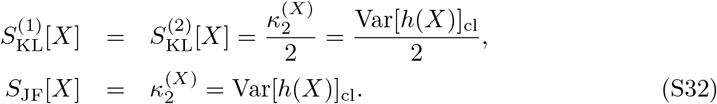

#### Analytical calculations of *K*_*D*_(*ξ*) and related relations given specific form of division count distributions

Here we derive (Eq. 22)-(Eq. 25) in the main text. We begin with the case where *Q*_cl_(*D*) follows a Poisson distribution. Let 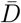 denote the chronological mean division count.

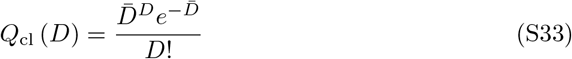

By the definition of *K*_*D*_(*ξ*),

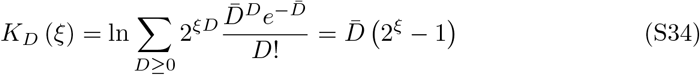

By the Taylor expansion of 2^*ξ*^ = *e*^*ξ* ln 2^, the *n*-th order cumulant is 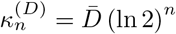.

Since

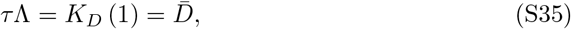

we derive

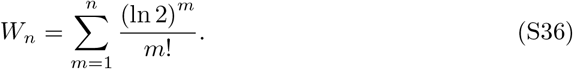

For example, 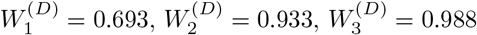, and 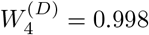. The first order derivative of *K*_*D*_(*ξ*) is

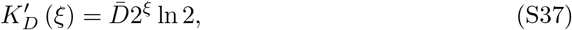

and thereby we have

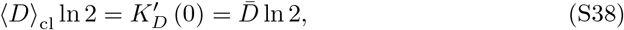

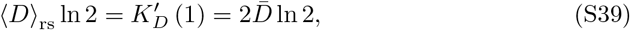

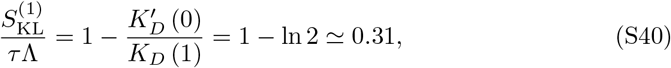

and

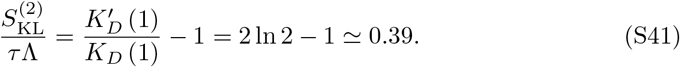

Next we derive *K*_*D*_(*ξ*) for binomial and negative binomial distributions. Let 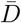 and 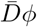 denote the mean and the variance of *Q*_cl_ (*D*). When *Q*_cl_ (*D*) is binomial,

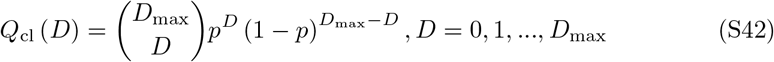

where *D*_max_ and *p* satisfy 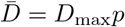 and 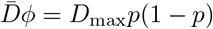; namely 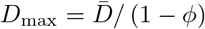 and *p* = 1 − *ϕ*. Therefore,

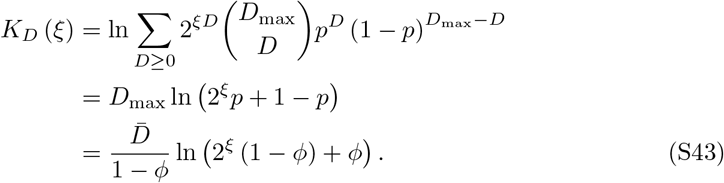

When *Q*_cl_ (*D*) is negative binomial,

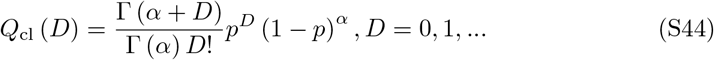

where *α* and *p* satisfy 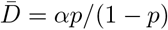 and 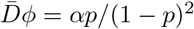; namely 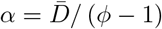 and *p* = 1 − *ϕ*^−1^. Therefore,

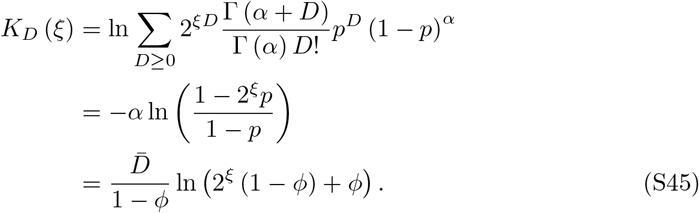

(Eq. S45) is exactly the same as (Eq. S43) as the function of 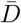, *ϕ*, and *ξ*. In addition, (Eq. S34) is the limiting form of (Eq. S43) and (Eq. S45) as *ϕ →* 1. Thus, (Eq. 22) in the main text represents *K*_*D*_(*ξ*) for Poisson, binomial or negative binomial *Q*_cl_(*D*).

The Taylor expansion of (Eq. 22) is obtained as follows:

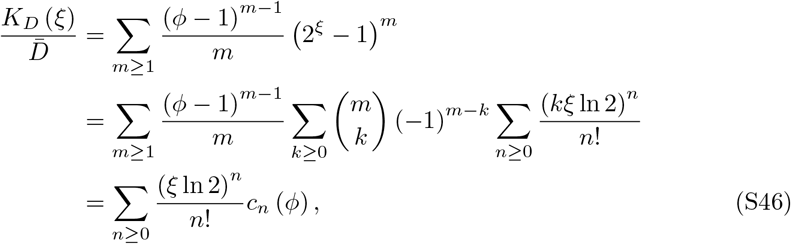

where

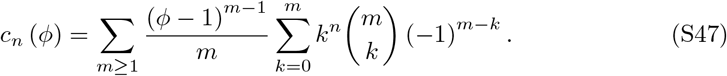

For the first five terms, for example, we have

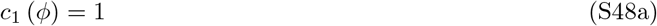

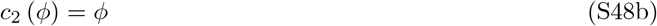

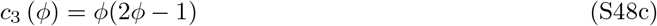

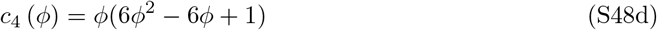

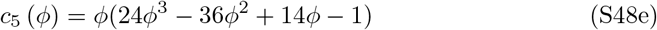

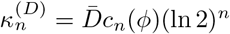 gives the *n*-th order cumulant.

The first order derivative of (Eq. 22) is

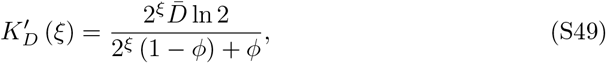

and thereby we obtain

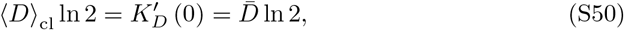

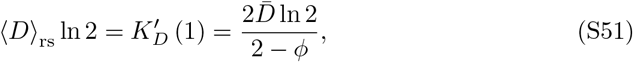

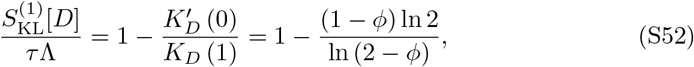

and

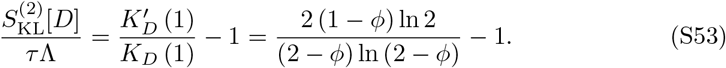

(Eq. S52) and (Eq. S53) equal if and only if

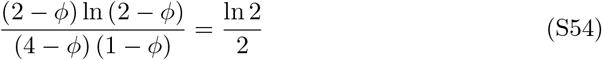

This equation has two roots *ϕ* = 0 and *ϕ* = *ϕ*_0_ = 0.5857… and LHS *>* RHS if and only if 0 *< ϕ < ϕ*_0_.

#### Long-term limit for gamma-distributed uncorrelated generations times

Let *g* (*x*) and *z* denote the probability density function of generation time *x* and the mean number of offsprings per generation, respectively. We assume that the generation time correlation between parent and offspring can be ignored; i.e., *g*(*x*) gives the probability density that offspring’s generation time becomes *x*. The Malthusian parameter *λ* is the real root of the so-called Euler-Lotka equation [40]:

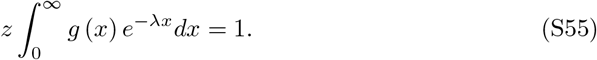

We remark that (Eq. S55) also holds for correlated generation times such as Markov models [41] by reinterpreting *g*(*x*) as the probability distribution of generation times of parent cells across a steadily growing population. In such cases, *g*(*x*) depends on *z*, and we cannot treat *g*(*x*) in (Eq. S61) independent of *z* = 2^*ξ*^. Here, we ignore any transgenerational correlations in generation time to illustrate the effect of the variation in generation time on *K*_*D*_(*ξ*) and selection strength measures with simple calculations. For this purpose, we further choose gamma distributions as *g*(*x*), i.e.,

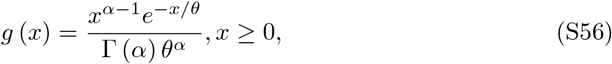

where *α >* 0 is a shape parameter; and *θ >* 0 is a scale parameter. In this case, the Malthusian parameter is

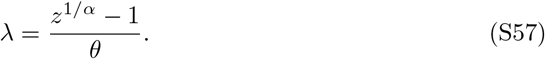

Using the mean and the variance of generation times (⟨*τ⟩*_*g*_ and *V ar* [*τ*]_*g*_, respectively), *α* and *θ* are expressed as

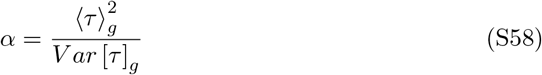

and

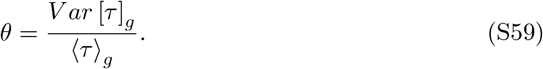

The probability distribution of division count *Q*_cl_(*D*), in this case, is known as gamma count distribution [42]. Though any closed-form expression of the corresponding cumulant generating function is not known, it has a simple limiting form for *τ →∞* as shown below. We define the rescaled cumulant generating function by

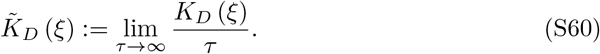

Since 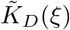 represents the population growth rate, or Malthusian parameter with the mean number of offspring *z* = 2^*ξ*^, we have

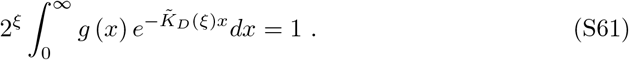

When *g* is a gamma distribution with a shape parameter *α* and a scale parameter *θ*, we obtain

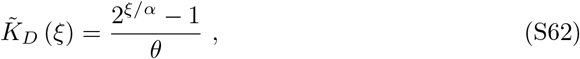

and

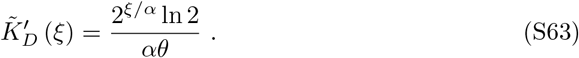

Note that *α* = 1 corresponds to the case where division counts follow the Poisson distribution with mean *θ*^−1^. The scaled key quantities derived from 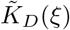 are as follows.

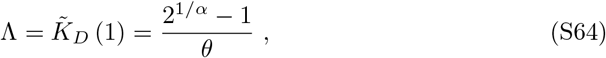

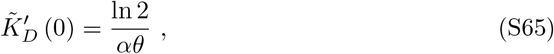

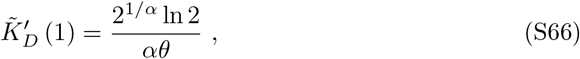

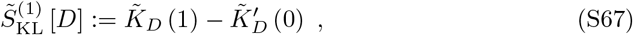

and

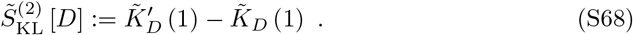

Hence,

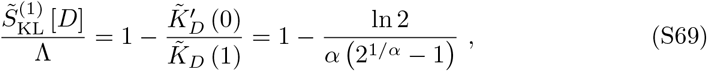

and

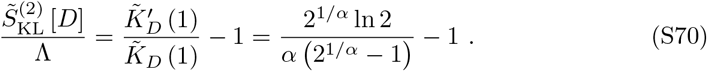

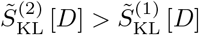 is always true for 0 *< α < ∞* because

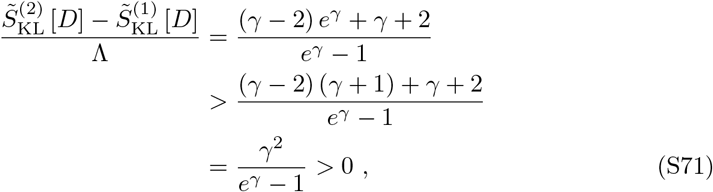

where *γ* = *α*^−1^ ln 2 and the inequality *e*^*γ*^ *>* 1 + *γ* (*γ >* 0) are used.

Since the Taylor expansion of 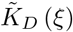 at *ξ* = 0 is

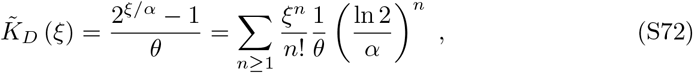

the time-scaled *n*-th order fitness cumulant is

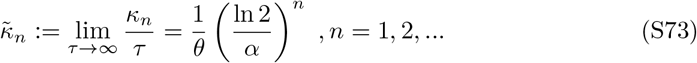

Therefore,

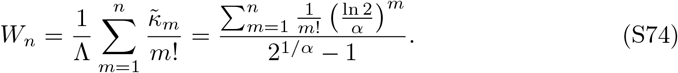

## Supplemental table

**Supplementary Table S1.** Summary of cellular species, culture conditions, and observation setup used in the experiments in Fig. 5.

| Species | Label | Strain | Medium | Temperature (°C) | Device | Source |
| --- | --- | --- | --- | --- | --- | --- |
| *E. coli* | rpoS-mcherry<br>glucose_30°C | MG1655 F3<br>rpoS-mcherry<br>/pUA66-PrpsL-gfp | M9 minimal medium<br>+0.2%(w/v) glucose<br>+1/2 MEM amino<br>acids solution (Sigma) | 30 | Microchamber<br>array | This study |
| *E. coli* | rpoS-mcherry<br>glucose_37°C | MG1655 F3<br>rpoS-mcherry<br>/pUA66-PrpsL-gfp | M9 minimal medium<br>+0.2%(w/v) glucose<br>+1/2 MEM amino<br>acids solution (Sigma) | 37 | Microchamber<br>array | This study |
| *E. coli* | rpoS-mcherry<br>glycerol_37°C | MG1655 F3<br>rpoS-mcherry<br>/pUA66-PrpsL-gfp | M9 minimal medium<br>+0.1%(v/v) glycerol<br>+1/2 MEM amino<br>acids solution (Sigma) | 37 | Microchamber<br>array | This study |
| *E. coli* | f3nw<br>-sm | F3NW | M9 minimal medium<br>+0.2%(w/v) glucose<br>+1/2 MEM amino<br>acids solution (Sigma)<br>+0.1 mM Isopropyl β-D-1<br>thiogalactopyranoside (IPTG) | 37 | Agar pad | [13] |
| *E. coli* | f3nw<br>+sm | F3NW | M9 minimal medium<br>+0.2%(w/v) glucose<br>+1/2 MEM amino<br>acids solution (Sigma)<br>+0.1 mM Isopropyl β-D-1<br>thiogalactopyranoside (IPTG)<br>+100 μg/ml streptomycin | 37 | Agar pad | [13] |
| *E. coli* | f3ptn001<br>-sm | F3/pTN001 | M9 minimal medium<br>+0.2%(w/v) glucose<br>+1/2 MEM amino<br>acids solution (Sigma)<br>+0.1 mM Isopropyl β-D-1<br>thiogalactopyranoside (IPTG) | 37 | Agar pad | [13] |
| *E. coli* | f3ptn001<br>+sm | F3/pTN001 | M9 minimal medium<br>+0.2%(w/v) glucose<br>+1/2 MEM amino<br>acids solution (Sigma)<br>+0.1 mM Isopropyl β-D-1<br>thiogalactopyranoside (IPTG)<br>+200 μg/ml streptomycin | 37 | Agar pad | [13] |
| *M. smegmatis* | mc<sup>2</sup>155<br>7H9 | mc<sup>2</sup>155 | Middlebrook 7H9 medium<br>+0.5% albumin +0.2% glucose<br>+0.085% NaCl +0.5% glycerol<br>+0.05% Tween-80 | 37 | Membrane<br>cover | [25] |
| *S. pombe* | EMM28 | HN0025 | Edinburgh minimal medium<br>+2% (w/v) glucose | 28 | Mother<br>machine | [26] |
| *S. pombe* | EMM30 | HN0025 | Edinburgh minimal medium<br>+2%(w/v) glucose | 30 | Mother<br>machine | [26] |
| *S. pombe* | EMM32 | HN0025 | Edinburgh minimal medium<br>+2%(w/v) glucose | 32 | Mother<br>machine | [26] |
| *S. pombe* | EMM34 | HN0025 | Edinburgh minimal medium<br>+2%(w/v) glucose | 34 | Mother<br>machine | [26] |
| *S. pombe* | YE28 | HN0025 | Yeast extract medium<br>+3%(w/v) glucose | 28 | Mother<br>machine | [26] |
| *S. pombe* | YE30 | HN0025 | Yeast extract medium<br>+3%(w/v) glucose | 30 | Mother<br>machine | [26] |
| *S. pombe* | YE34 | HN0025 | Yeast extract medium<br>+3%(w/v) glucose | 34 | Mother<br>machine | [26] |
| L1210<br>mouse leukemia<br>cell | L1210<br>RPMI-1640 | L1210<br>(ATCC CCL-219) | RPMI-1640 medium (Wako)<br>+10% fetal bovine serum (Biosera)<br>under 5% CO<sub>2</sub> atmosphere | 37 | Mother<br>machine | [22] |

**Supplementary Table S2.**
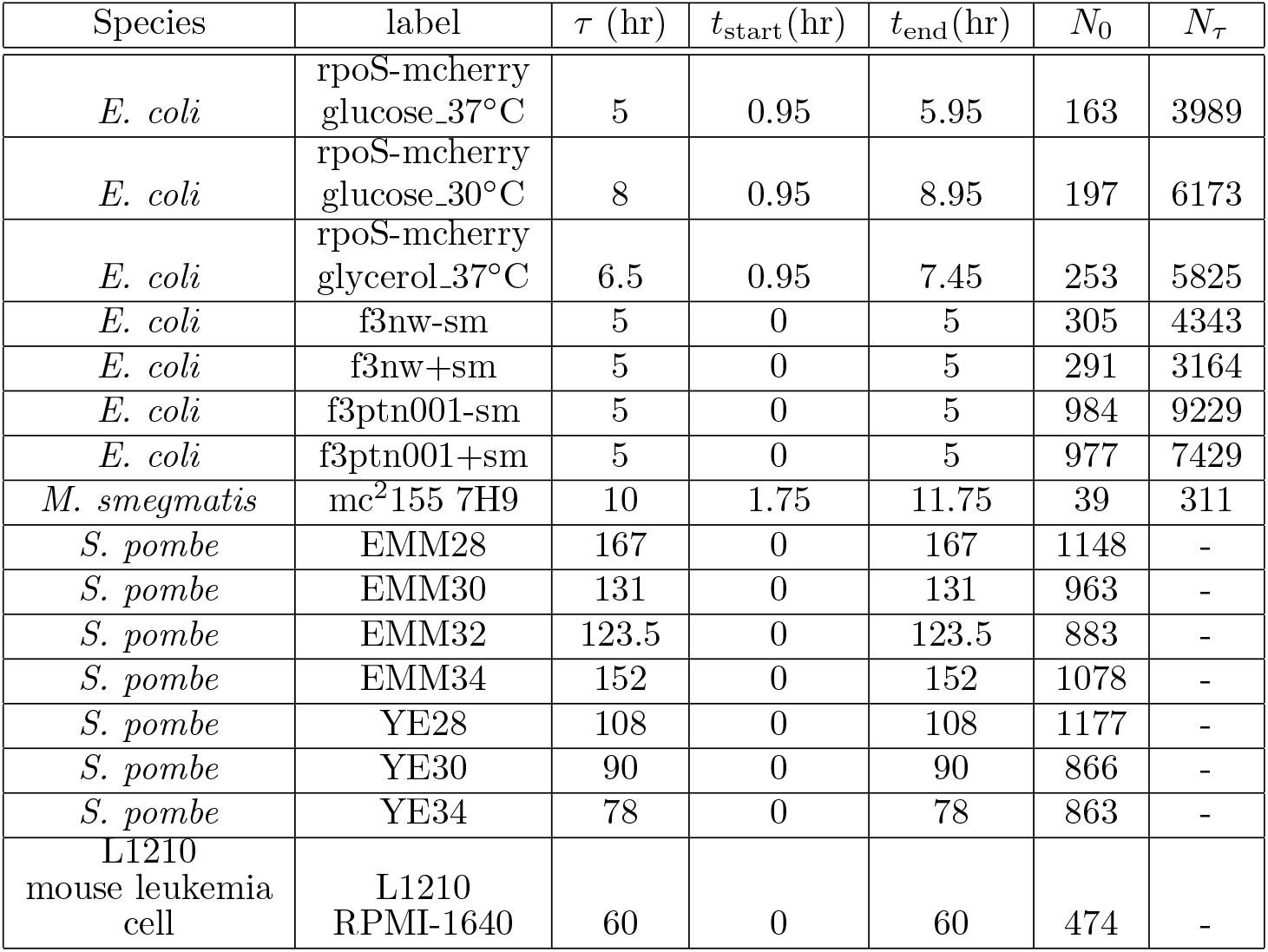
Summary of the data used in the analysis in Fig. 5. *t*_start_ and *t*_end_ are the start and end times for the analysis time window *τ*.

| Species | label | $\tau$ (hr) | $t_{\text{start}}$(hr) | $t_{\text{end}}$(hr) | $N_0$ | $N_\tau$ |
| --- | --- | --- | --- | --- | --- | --- |
| *E. coli* | rpoS-mcherry<br>glucose.37°C | 5 | 0.95 | 5.95 | 163 | 3989 |
| *E. coli* | rpoS-mcherry<br>glucose.30°C | 8 | 0.95 | 8.95 | 197 | 6173 |
| *E. coli* | rpoS-mcherry<br>glycerol.37°C | 6.5 | 0.95 | 7.45 | 253 | 5825 |
| *E. coli* | f3nw-sm | 5 | 0 | 5 | 305 | 4343 |
| *E. coli* | f3nw+sm | 5 | 0 | 5 | 291 | 3164 |
| *E. coli* | f3ptn001-sm | 5 | 0 | 5 | 984 | 9229 |
| *E. coli* | f3ptn001+sm | 5 | 0 | 5 | 977 | 7429 |
| *M. smegmatis* | mc<sup>2</sup>155 7H9 | 10 | 1.75 | 11.75 | 39 | 311 |
| *S. pombe* | EMM28 | 167 | 0 | 167 | 1148 | - |
| *S. pombe* | EMM30 | 131 | 0 | 131 | 963 | - |
| *S. pombe* | EMM32 | 123.5 | 0 | 123.5 | 883 | - |
| *S. pombe* | EMM34 | 152 | 0 | 152 | 1078 | - |
| *S. pombe* | YE28 | 108 | 0 | 108 | 1177 | - |
| *S. pombe* | YE30 | 90 | 0 | 90 | 866 | - |
| *S. pombe* | YE34 | 78 | 0 | 78 | 863 | - |
| L1210<br>mouse leukemia<br>cell | L1210<br>RPMI-1640 | 60 | 0 | 60 | 474 | - |

## Supplemental figures

**Figure S1.**
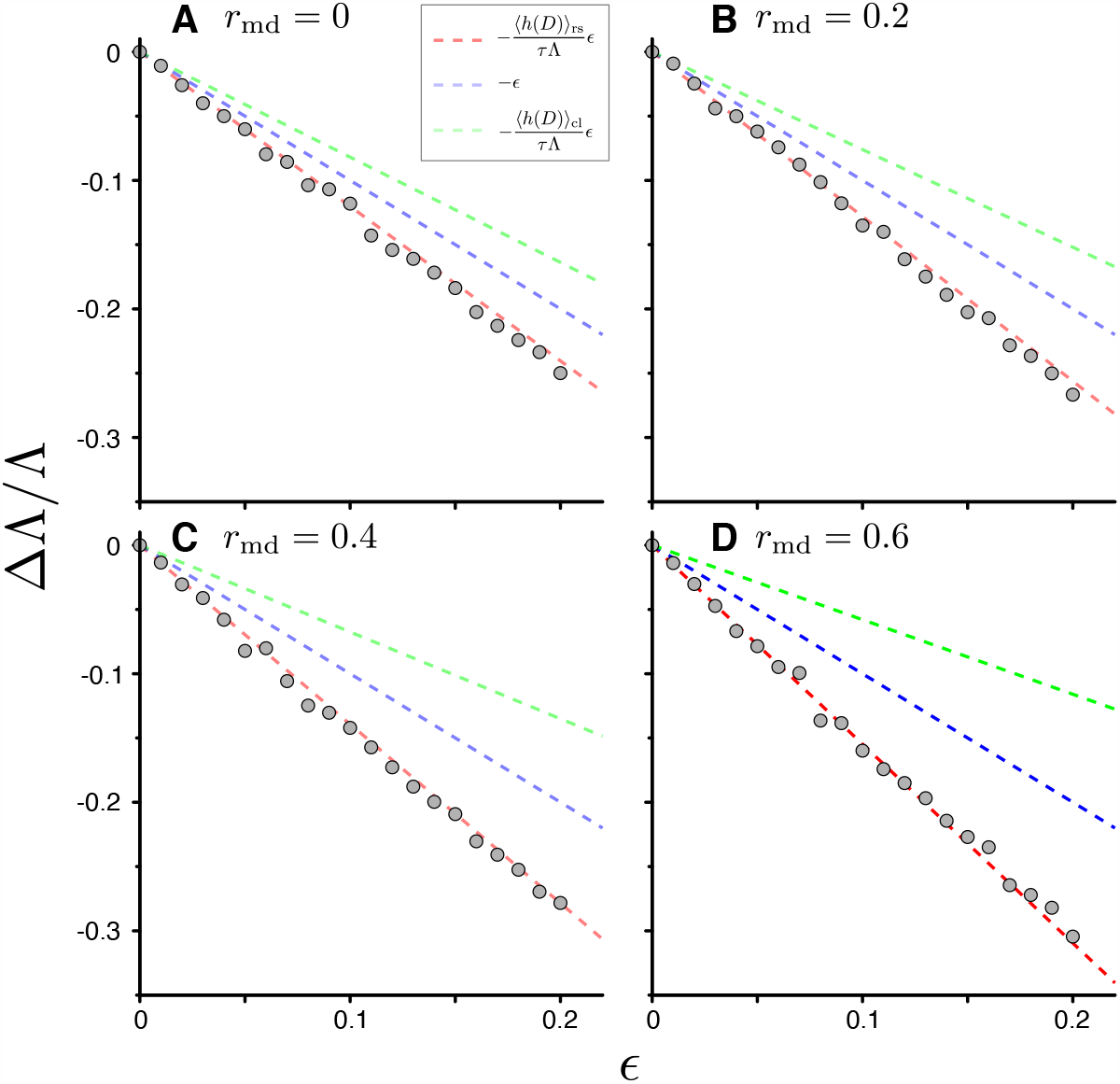
Response of population growth rate to cell removal perturbation with positive mother-daughter correlations of generation time. We conducted the simulations of cell population growth with positive mother-daughter correlations (Gray points). We considered the generation time distribution that follows a gamma distribution with shape parameter 2 (*g*_2_(*x*) in Fig. 4B). We generated correlated generation time of individual cells based on the mother cell’s value with the algorithm in Materials and Methods. We randomly removed the individual cells from the population with the probability 1 − 2^−*ϵ*^ after each cell division. The broken red lines represent the theoretical prediction of the changes 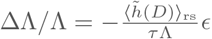. The lines of ΔΛ*/*Λ = −*ϵ* (blue) and 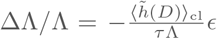 (green) were also shown for reference. **A**. The result with no mother-daughter correlation of generation time (*r*_md_ = 0). **B**. *r*_md_ = 0.2. **C**. *r*_md_ = 0.4. **D**. *r*_md_ = 0.6.

**Figure S2.**
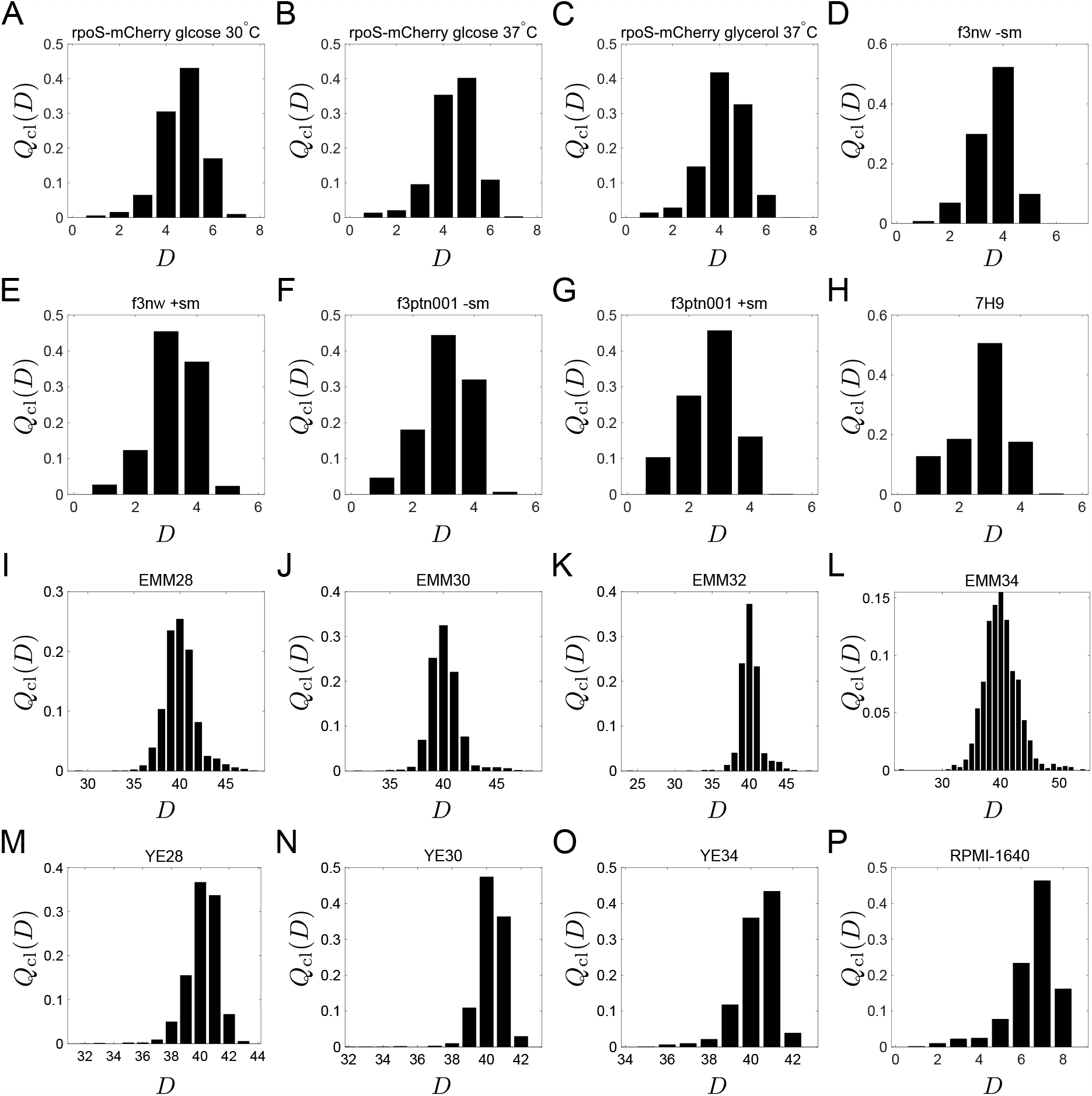
Chronological distributions of division count, *Q*_cl_(*D*). **A-G**. *E. coli*. **H**. *M. smegmatis*. **I-O**. *S. pombe*. **P**. L1210 mouse leukemia cells.

**Figure S3.**
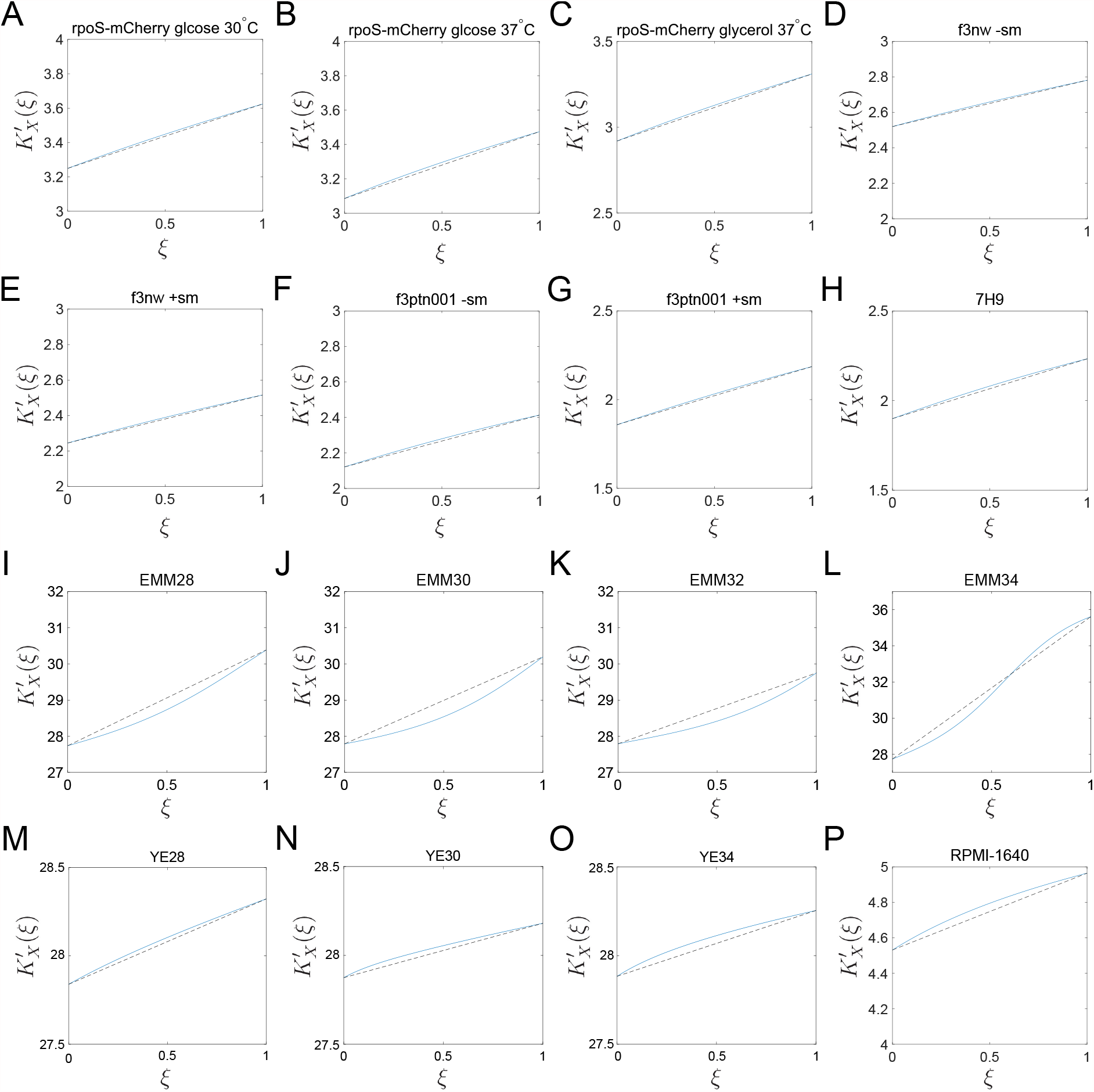
Graphical representation of 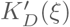. A-G. *E. coli*. H. *M. smegmatis*. I-O. *S. pombe*. P. L1210 mouse leukemia cells.

## References

1. Balázsi G, van Oudenaarden A, Collins JJ. Cellular decision making and biological noise: from microbes to mammals. Cell. 2011;144(6):910–25. doi:10.1016/j.cell.2011.01.030.

2. Elowitz MB, Levine AJ, Siggia ED, Swain PS. Stochastic gene expression in a single cell. Science. 2002;297(5584):1183–6. doi:10.1126/science.1070919.

3. Kelly CD, Rahn O. The Growth Rate of Individual Bacterial Cells. Journal of Bacteriology. 1932;23(2):147–53.

4. Powell EO. Growth rate and generation time of bacteria, with special reference to continuous culture. Journal of General Microbiology. 1956;15(3):492–511.

5. Wakamoto Y, Ramsden J, Yasuda K. Single-cell growth and division dynamics showing epigenetic correlations. Analyst. 2005;130(3). doi:10.1039/b409860a.

6. Wang P, Robert L, Pelletier J, Dang WL, Taddei F, Wright A, et al. Robust growth of Escherichia coli. Current Biology. 2010;20(12):1099–1103. doi:10.1016/j.cub.2010.04.045.

7. Cerulus B, New AM, Pougach K, Verstrepen KJ. Noise and epigenetic inheritance of single-cell division times influence population fitness. Current Biology. 2016;26(9):1138–1147. doi:10.1016/j.cub.2016.03.010.

8. Susman L, Kohram M, Vashistha H, Nechleba JT, Salman H, Brenner N. Individuality and slow dynamics in bacterial growth homeostasis. Proceedings of the National Academy of Sciences. 2018;115(25):E5679–E5687. doi:10.1073/pnas.1615526115.

9. Leibler S, Kussell E. Individual histories and selection in heterogeneous populations. Proceedings of the National Academy of Sciences of the United States of America. 2010;107(29):13183–8. doi:10.1073/pnas.0912538107.

10. Hashimoto M, Nozoe T, Nakaoka H, Okura R, Akiyoshi S, Kaneko K, et al. Noise-driven growth rate gain in clonal cellular populations. Proceedings of the National Academy of Sciences. 2016;113(12):3251–3256. doi:10.1073/pnas.1519412113.

11. Rochman ND, Popescu DM, Sun SX. Ergodicity, hidden bias and the growth rate gain. Physical Biology. 2018;15(3). doi:10.1088/1478-3975/aab0e6.

12. Stewart EJ, Madden R, Paul G, Taddei F. Aging and death in an organism that reproduces by morphologically symmetric division. PLoS Biology. 2005;3:e45. doi:10.1371/journal.pbio.0030045.

13. Nozoe T, Kussell E, Wakamoto Y. Inferring fitness landscapes and selection on phenotypic states from single-cell genealogical data. PLoS Genetics. 2017;13(3):e1006653. doi:10.1371/journal.pgen.1006653.

14. García-Garćia R, Genthon A, Lacoste D. Linking lineage and population observables in biological branching processes. Physical Review E. 2019;99(4):1–12. doi:10.1103/PhysRevE.99.042413.

15. Levien E, GrandPre T, Amir A. Large Deviation Principle Linking Lineage Statistics to Fitness in Microbial Populations. Physical Review Letters. 2020;125(4):048102. doi:10.1103/PhysRevLett.125.048102.

16. Genthon A, Lacoste D. Fluctuation relations and fitness landscapes of growing cell populations. Scientific Reports. 2020;10(1):1–13. doi:10.1038/s41598-020-68444-x.

17. Wakamoto Y, Grosberg AY, Kussell E. Optimal lineage principle for age-structured populations. Evolution. 2012;66(1):115–34. doi:10.1111/j.1558-5646.2011.01418.x.

18. Kobayashi TJ, Sughiyama Y. Fluctuation Relations of Fitness and Information in Population Dynamics. Physical Review Letters. 2015;115:238102. doi:10.1103/PhysRevLett.115.238102.

19. Thomas P. Single-cell histories in growing populations: relating physiological variability to population growth. bioRxiv. 2017;doi:10.1101/100495.

20. Lin J, Amir A. The effects of stochasticity at the single-cell level and cell size control on the population growth. Cell systems. 2017;5:358–367.e4. doi:10.1016/j.cels.2017.08.015.

21. Nozoe T, Kussell E. Cell Cycle Heritability and Localization Phase Transition in Growing Populations. Physical Review Letters. 2020;125(26):268103. doi:10.1103/PhysRevLett.125.268103.

22. Seita A, Nakaoka H, Okura R, Wakamoto Y. Intrinsic growth heterogeneity of mouse leukemia cells underlies differential susceptibility to a growth-inhibiting anticancer drug. PLOS ONE. 2021;16(2):e0236534.

23. Mosheiff N, Martins BMC, Pearl-Mizrahi S, Grünberger A, Helfrich S, Mihalcescu I, et al. Inheritance of cell-cycle duration in the presence of periodic forcing. Physical Review X. 2018;8(2):021035. doi:10.1103/PhysRevX.8.021035.

24. Kuchen EE, Becker NB, Claudino N, Höfer T. Hidden long-range memories of growth and cycle speed correlate cell cycles in lineage trees. eLife. 2020;9:e51002. doi:10.7554/eLife.51002.

25. Wakamoto Y, Dhar N, Chait R, Schneider K, Signorino-Gelo F, Leibler S, et al. Dynamic persistence of antibiotic-stressed mycobacteria. Science. 2013;339:91–5. doi:10.1126/science.1229858.

26. Nakaoka H, Wakamoto Y. Aging, mortality, and the fast growth trade-off of Schizosaccharomyces pombe. PLOS Biology. 2017;15(6):e2001109. doi:https://doi.org/10.1371/journal.pbio.2001109.

27. Inoue I, Wakamoto Y, Moriguchi H, Okano K, Yasuda K. On-chip culture system for observation of isolated individual cells. Lab on a chip. 2001;1(1):50–5. doi:10.1039/b103931h.

28. Battesti A, Majdalani N, Gottesman S. The RpoS-mediated general stress response in Escherichia coli. Ann Rev Microbiol. 2011;34:189–213. doi:10.1146/annurev-micro-090110-102946.

29. Jun S, Si F, Pugatch R, Scott M. Fundamental principles in bacterial physiology—history, recent progress, and the future with focus on cell size control: a review. Reports on Progress in Physics. 2018;81(5):056601. doi:10.1088/1361-6633/aaa628.

30. Kohram M, Vashistha H, Leibler S, Xue B, Salman H. Bacterial Growth Control Mechanisms Inferred from Multivariate Statistical Analysis of Single-Cell Measurements. Current Biology. 2021;31(5):955–964.e4. doi:https://doi.org/10.1016/j.cub.2020.11.063.

31. Lambert G, Kussell E. Memory and Fitness Optimization of Bacteria under Fluctuating Environments. PLoS genetics. 2014;10(9):e1004556. doi:10.1371/journal.pgen.1004556.

32. Kussell E, Leibler S. Phenotypic diversity, population growth, and information in fluctuating environments. Science. 2005;309:2075–8. doi:10.1126/science.1114383.

33. Quinn JJ, Jones MG, Okimoto RA, Nanjo S, Chan MM, Yosef N, et al. Single-cell lineages reveal the rates, routes, and drivers of metastasis in cancer xenografts. Science. 2021;371(6532). doi:10.1126/science.abc1944.

34. Filipczyk A, Marr C, Hastreiter S, Feigelman J, Schwarzfischer M, Hoppe PS, et al. Network plasticity of pluripotency transcription factors in embryonic stem cells. Nature Cell Biology. 2015;17:1235–1246. doi:10.1038/ncb3237.

35. Frieda KL, Linton JM, Hormoz S, Choi J, Chow KHK, Singer ZS, et al. Synthetic recording and in situ readout of lineage information in single cells. Nature. 2017;541:107–111. doi:10.1038/nature20777.

36. Chow KHK, Budde MW, Granados AA, Cabrera M, Yoon S, Cho S, et al. Imaging cell lineage with a synthetic digital recording system. Science. 2021;372. doi:10.1126/science.abb3099.

37. Zaslaver A, Bren A, Ronen M, Itzkovitz S, Kikoin I, Shavit S, et al. A comprehensive library of fluorescent transcriptional reporters for Escherichia coli. Nature Methods. 2006;3:623–8. doi:10.1038/nmeth895.

38. Edelstein AD, Tsuchida MA, Amodaj N, Pinkard H, Vale RD, Stuurman N. Advanced methods of microscope control using μ Manager software. Journal of Biological Methods. 2014;1(2):10. doi:10.14440/jbm.2014.36.

39. Schneider CA, Rasband WS, Eliceiri KW. NIH Image to ImageJ: 25 years of image analysis. Nature Methods. 2012;9(7):671–5. doi:10.1038/nmeth.2089.

40. Fisher RA. The Genetical Theory of Natural Selection. Oxford University Press; 1930.

41. Lebowitz JL, Rubinow SI. A theory of the age and generation time distribution of a microbial population. Journal of Mathematical Biology. 1974;1:17–36.

42. Winkelmann R. Duration Dependence and Dispersion in Count-Data Models. Journal of Business & Economic Statistics. 1995;13(4):467–474. doi:10.1080/07350015.1995.10524620.

